# Dynamic DNA methylation reveals novel *cis-*regulatory elements in murine hematopoiesis

**DOI:** 10.1101/2022.06.02.493896

**Authors:** Maximilian Schönung, Mark Hartmann, Stephen Krämer, Sina Stäble, Mariam Hakobyan, Emely Kleinert, Theo Aurich, Defne Cobanoglu, Florian H. Heidel, Stefan Fröhling, Michael D. Milsom, Matthias Schlesner, Pavlo Lutsik, Daniel B. Lipka

**Author notes:** Correspondence to: Daniel B. Lipka, Section Translational Cancer Epigenomics, Division Translational Medical Oncology, German Cancer Research Center (DKFZ) & National Center for Tumor Diseases (NCT), Heidelberg, Germany.

## Abstract

**Background:** The differentiation of hematopoietic stem and progenitor cells (HSPCs) to terminally differentiated immune cells is accompanied by large-scale remodeling of the DNA methylation landscape. While significant insights into the molecular mechanisms of hematopoietic tissue regeneration were derived from mouse models, profiling of DNA methylation changes has been hampered by high cost or low resolution using the methods available. This problem has been overcome by the recent development of the Infinium Mouse Methylation BeadChip (MMBC) array, facilitating methylation profiling of the mouse genome at single CpG resolution at affordable cost.

**Results:** We extended the RnBeads package to provide a computational framework for the analysis of MMBC data. This framework was applied to a newly generated MMBC reference map of mouse hematopoiesis encompassing nine different cell types. The analysis of dynamically regulated CpG sites showed progressive and unidirectional DNA methylation changes from HSPCs to differentiated hematopoietic cells and allowed the identification of lineage- and cell type-specific DNA methylation programs. Comparison to previously published catalogues of cis-regulatory elements (CREs) revealed 12,856 novel putative CREs which were dynamically regulated by DNA methylation (mdCREs). These mdCREs were predominantly associated with patterns of cell type-specific DNA hypomethylation and could be identified as epigenetic control regions regulating the expression of key hematopoietic genes during differentiation.

**Conclusions:** We established a publicly available analysis pipeline for MMBC datasets and provide a DNA methylation atlas of mouse hematopoiesis. This resource allowed us to identify novel putative CREs involved in hematopoiesis and will serve as a platform to study epigenetic regulation of normal and malignant hematopoiesis.

## BACKGROUND

The hematopoietic system of the mouse is among the best studied model systems for regenerative tissues. Over the last decades, a roadmap of murine hematopoietic differentiation has been generated whereby hematopoietic stem and progenitor cells (HSPCs) reside on the top of a hierarchically organized system. These cells give rise to lineage-committed progenitor cells, which then differentiate into mature hematopoietic cells. This differentiation process is tightly regulated in order to ensure faithful production of hematopoietic cells according to the actual needs of the organism. Epigenetic regulation plays a key role in these processes and DNA methylation has emerged as an indispensable epigenetic mark required for maintenance of hematopoietic stem cell (HSC) function and for faithful hematopoietic differentiation [1–4]. DNA methylation refers to the covalent attachment of a methyl-group to cytosine residues in the DNA, which in mammals mainly occurs in a CpG sequence context [5]. This process is catalyzed by three enzymes of the DNA methyltransferase (DNMT) family, namely DNMT1, DNMT3A and DNMT3B. The latter two are required for the *de novo* establishment of DNA methylation marks whereas DNMT1 is a maintenance DNA methyltransferase, which propagates DNA methylation patterns following DNA replication [6].

The observation that DNMT3A is among the most frequently mutated genes in human acute myeloid leukemia (AML) [7, 8] led to a variety of studies investigating the role of DNA methylation in healthy and malignant hematopoiesis [9–15]. An initial study using a custom array platform revealed global DNA methylation plasticity during murine progenitor differentiation and observed a global hypermethylation accompanying lymphoid lineage commitment [10]. This work was further refined using reduced representation bisulfite sequencing (RRBS) which assessed DNA methylation patterns at single CpG resolution. The authors demonstrated that myeloid TF binding sites are methylated during lymphoid differentiation, suggesting a suppression of opposing lineage identities [9]. An integrated genome-wide DNA methylome and transcriptome map of the HSPC compartment subsequently identified a set of genes whose expression is at least in part regulated by DNA methylation programming and provided further support for the importance of epigenetic programming in normal hematopoiesis [11, 12]. Since an in-depth characterization of regulatory sites is required to achieve a comprehensive understanding of hematopoietic differentiation processes at the systems level, the robust identification and functional annotation of *cis-*regulatory elements (CREs) via computational epigenomics is a critically important task. Many consortia, like *The Immunological Genome Project* (ImmGen) or the *Validated Systematic Integration of Hematopoietic Epigenomes* (VISION), have intensively worked on mapping and stratifying CREs which are dynamically regulated during hematopoiesis [16, 17]. However, these CRE catalogues are solely defined by open chromatin and histone modification patterns and thus lack a comprehensive annotation of dynamic DNA methylation changes in mouse hematopoiesis. In contrast, the *Encyclopedia of DNA Elements* (ENCODE) analyzed DNA methylation, histone marks, CTFTC and DNaseI binding across multiple organs and developmental stages during mouse ontogenesis [18]. Nevertheless, this consortium did not specifically analyze cell types of the hematopoietic system and their registry of candidate CREs (cCREs) is based on H3K4me3, H3K27ac, CTCF and DNaseI, and does not include DNA methylation.

The lack of a systematic DNA methylation layer in current CRE atlases might in part be explained by lack of robust, reproducible and cost-efficient methods for DNA methylation analysis in mouse cells. Genome-wide methods include *whole-genome bisulfite sequencing* (WGBS) and *reduced representation bisulfite sequencing* (RRBS). While RRBS can be performed at relatively low costs, this method provides only limited information on the DNA methylome with a strong bias towards CpG-dense regions. In contrast, WGBS provides unbiased information covering all CpGs throughout the genome. However, this is associated with comparatively high costs, resulting from the requirement to sequence with a sufficient genome coverage to accurately estimate the DNA methylation level of individual CpGs. As a result, WGBS is not amenable for large-scale DNA methylation analysis of mouse models.

In human studies, these limitations were overcome by the introduction of Infinium DNA Methylation Bead Chip arrays, which in their latest version allow the highly reproducible analysis of up to 850,000 CpG-sites at low cost, facilitating the study of large patient cohorts [19–21]. However, until recently, this technology was not available for mouse samples. This changed with the recent introduction of the Infinium Mouse Methylation BeadChip (MMBC) array in 2020, which now allows DNA methylome analyses in mice at affordable cost. The MMBC array interrogates 285,000 CpG sites across the mouse genome, including promoter regions of more than 28,000 protein-coding transcripts and 60,000 enhancers. However, meaningful data analysis is likely to still pose a significant hurdle for many groups who do not have specialized bioinformatic support in this area.

We established an easy-to-use computational pipeline for the analysis of MMBC data that is based on the RnBeads Bioconductor package [22, 23]. As a proof-of-concept, we applied this pipeline to MMBC data generated from nine hematopoietic cell types across all major lineages to provide an atlas of DNA methylation patterns of the murine hematopoietic system, which can serve as a resource for the scientific community. In-depth analysis of this data set identified DNA methylation programs associated with hematopoietic lineage commitment and revealed novel candidate CREs which likely contribute to differentiation and regulation processes in hematopoiesis.

## RESULTS

### Implementation of an MMBC-array analysis framework in RnBeads

To facilitate seamless analysis of MMBC-array data, we extended our earlier published R-package RnBeads and used it to build a comprehensive computational workflow for the present study [22, 23]. First, we converted the manifest file provided by the manufacturer into an RnBeads-compatible MMBC-array annotation, which we added to the RnBeads.mm10 companion data package (see Methods for further details). Second, we adapted and generalized the core functionality of RnBeads to support the new array type. As a result, our RnBeads-based computational workflow allows for an automated and user-friendly processing and analysis of MMBC data, which is in full equivalence with the RnBeads workflows for 450k/EPIC-arrays. In particular, data loading and quality control of the unprocessed IDAT files can be executed using a single R function with global pipeline configuration options. The subsequent pre-processing module features several methods for background correction, automated filtering and normalization. The majority of QC and preprocessing procedures already available in RnBeads for human 450k/EPIC arrays also work for MMBC-array data, including: control probe visualization; negative control- and out-of-band-based background subtraction; multiplicative dye bias correction; and most probe filtering steps. Additional downstream analysis modules allow users to perform standard exploratory analysis, such as dimensionality reduction; inference of batch effects and major covariates; calling of differentially methylated probes (DMPs); and execution of reference-based cell type deconvolution. Each of the modules generates an interactive HTML-report for documentation and reproducibility purposes (**Figure 1A**). The updated package, featuring MMBC support starting from version 2.9.4, is freely available on Bioconductor [24]. Furthermore, we generated a comprehensive functional annotation of the MMBC array including mapping of probes to regulatory regions in the mouse genome (mm10) and gene annotations (**Supplementary Table 1**). Finally, we established a bioinformatic workflow for *de novo* identification of methylation dynamic CREs (mdCREs) throughout hematopoietic differentiation. Integration with published gene expression data allowed us to propose putative functional gene annotations for a subset of these novel mdCREs (**Figure 1A**).

**Figure 1.**
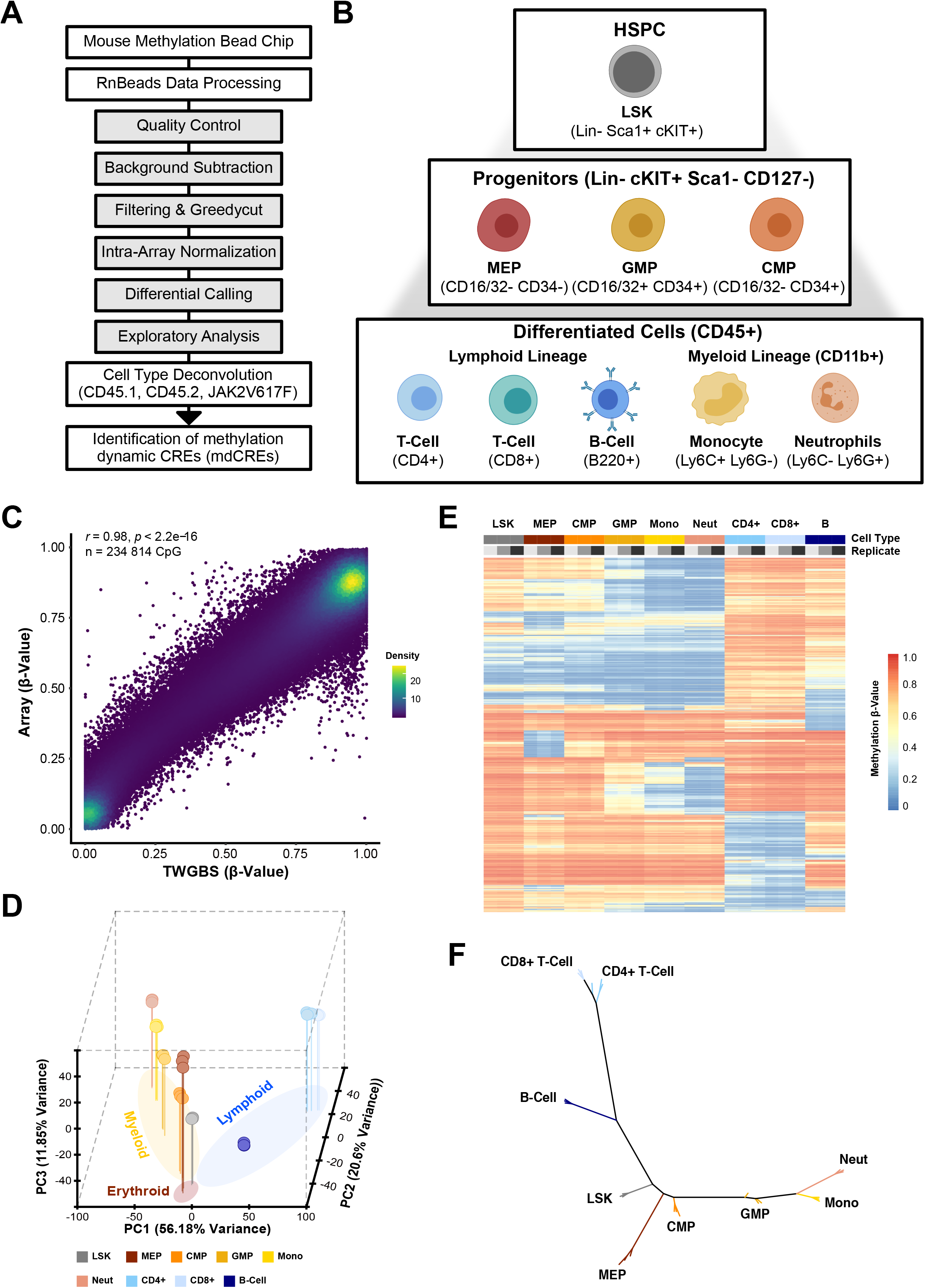
Mouse methylation bead chip reference map of hematopoietic differentiation. **(A)** Schematic depicting the computational pipeline that was developed and implemented within the RnBeads Bioconductor package. This pipeline includes quality control, pre-processing (i.e. background subtraction, filtering, normalization), differential methylation calling, cell type deconvolution and *de novo* identification of methylation dynamic CREs, and visualization of the mouse methylation bead chip data. **(B)** Overview on the hematopoietic cell types isolated by flow cytometry and analyzed in the present study. **(C)** Density dot plot showing the correlation of DNA methylation β-values derived from tagmentation-based whole genome bisulfite sequencing (TWGBS) and MMBC data of Lin-Sca1+ Kit+ (LSK) cells. All CpG sites with >10 reads in at least 2 TWGBS samples were included in this analysis (n=234,814 CpGs). Pearson correlation has been calculated. **(D)** Three-dimensional principal component analysis using the 5000 most variable CpG (mvCpG) sites separated major hematopoietic cell types and differentiation branches. The axes depict the principal components (PC) 1-3. **(E)** Heatmap depicting the hierarchical clustering of the 5000 mvCpGs. Columns represent the cell types and the biological replicates and the rows represent the CpGs. Rows were clustered using Manhattan distance and complete linkage. Columns were ordered based on known lineage relationships in the hematopoietic system. **(F)** Phylogenetic tree depicting the relationship of all cell types including the replicate information over three replicates for each cell type. The phylogenetic tree was calculated based on the methylation levels of the 5000 mvCpGs using the Manhattan distance metric.

### Mouse methylation bead chip reference map of murine hematopoietic differentiation

To study the DNA methylation patterns across the mouse hematopoietic system, nine hematopoietic cell types were prospectively isolated in biological triplicates from bone marrow of C57BL/6J mice by fluorescence-activated cell sorting (FACS) using a set of established cell surface markers (**Figure 1B**, **Supplementary Figure 1, Supplementary Table 2**). Genomic DNA was purified and analyzed on the MMBC platform, which allows the simultaneous analysis of the DNA methylation status of 285,000 CpGs throughout the mouse genome. The data was processed using our newly established RnBeads MMBC analysis pipeline. We then assessed the quality of the generated MMBC dataset by 1) evaluation of the internal control probe intensities for each step of the procedure with respect to their expected values; 2) interrogating the median Infinium bead count value and proportion of probes with low Infinium bead counts; and 3) assessing the proportion of probes with high detection p-values. Our QC assessment confirmed the overall high quality of the data generated with the MMBC array. Internal control probes demonstrated expected intensity ranges according to the annotation provided by the manufacturer. The median bead count was 24 [range 0 to 204] and all samples had more than 261,550 (99.98 %) probes with Infinium bead count >3. The number of low-quality probes with detection p-values > 0.001 ranged between 3 and 22 probes across all samples (**Supplementary Figure 2A**). Per CpG comparison of beta-values between biological replicates showed high correlation for all cell types as demonstrated for LSK (Pearson correlation, r = 0.998, p < 2.2e-16), further confirming the technical reproducibility (**Supplementary Figure 2B, C**). Additionally, inspection of methylation β-values before and after intra-array normalization revealed a minor shift of the underlying methylation values, indicating a uniformly high quality of the dataset (**Supplementary Figure 2D**). To further assess the quality of the MMBC data, we compared this with previously published data from analogous cell populations generated using either tagmentation-based whole-genome bisulfite sequencing (TWGBS), or reduced representation bisulfite sequencing (RRBS) data. The TWGBS dataset covered ∼2.48 x 10^5^ CpGs (100%) of the 2.48 x 10^5^ autosomal CpGs present on the MMBC array, while the RRBS dataset covered only ∼2.1 x 10^4^ (9%) of the autosomal CpGs represented on the MMBC array (**Supplementary Table 3**). After quality control and coverage filtering, 2.35 x 10^5^ CpGs (95% of autosomal MMBC CpGs) and 0.6 - 1.6 x 10^4^ CpGs (2.5 – 6.7% of autosomal MMBC CpGs) remained from the TWGBS and the RRBS datasets. Further analysis revealed a high correlation of the MMBC data with the published DNA methylation data sets, demonstrating the high quality of our MMBC data set (r = 0.98 for TWGBS and r = 0.97 - 0.98 for RRBS; **Figure 1C, Supplementary Figure 3**). Unsupervised analysis of DNA methylation patterns in the represented cell types using principal component analysis (PCA) showed that replicates from the same cell type cluster closely together. Thus, different cell types can be distinguished based on global DNA methylation differences whereby the lymphoid and myeloid lineages showed the most divergent DNA methylation patterns (**Figure 1D, E**). Phylogenetic tree analysis recapitulated the early branching of the lymphoid and the myeloid arms starting from immature LSK cells (**Figure 1F**). Furthermore, this analysis demonstrated that MEPs branch early from the myeloid arm, which is in line with a previously described erythroid/megakaryocytic-priming of CD55+ MPPs and CMPs [25]. In summary, we have generated a high quality MMBC reference data set of murine hematopoiesis that may serve as a resource for researchers working with mouse models spanning a wide spectrum of diseases.

### Identification of reference DNA methylation programs allows reliable cell type deconvolution from bulk samples

Previous studies of reference epigenomes have revealed cell type-specific DNA methylation patterns and underlined their suitability to infer the cellular composition of bulk samples [26]. Based on this knowledge, we tested if our MMBC data set could be used to perform reference-based cell type deconvolution of murine bone marrow samples. Therefore, cell type-specific differentially methylated probes (ctDMPs) were identified using a stringent filtering strategy. We selected CpGs which exhibited strong cell type-specific hypomethylation while at the same time maintaining uniformly high methylation levels across all other cell types (**Figure 2A; Supplementary Table 4**). Data from HSPCs and myeloid progenitor cells (CMPs and GMPs) were not included in this analysis, as these cell populations constitute only a minor fraction of cells in total bone marrow samples. Methylation data from CD4+ and CD8+ T cells were combined and considered as “T cells”, as our filtering strategy did not identify CpGs that would allow us to distinguish between these T-cell subsets. In total, this analysis identified a set of 201 ctDMPs which were used as features for cell type deconvolution (**Figure 2A, B**). Four different cell type deconvolution algorithms, which had previously been shown to work on human DNA methylation array data, were applied to compare their performance on the selected ctDMPs: the reference-based constrained projection (CP) algorithm (“Houseman”); the EpiDISH Cibersort (CBS) algorithm; the EpiDISH robust partial correlations (RPC) algorithm; and the reference-free constrained non-negative matrix factorization algorithm MeDeCom [27–30].The performance of these algorithms was tested on array data generated from total bone marrow samples isolated from two C57BL/6 wild type mice carrying either the Cd45.1 or the Cd45.2 allele, as well as from two bone marrow samples isolated from a myeloproliferative neoplasia mouse model. These mice expressed a conditional JAK2V617F knock-in allele which previously caused a shifted in the myeloid and erythroid compartment, thus allowing us to test the capability of the algorithms to predict disease-specific changes in the cell type composition. Flow cytometry-based measurements of all animals were used as a comparator data (**Figure 2C**). The mean absolute error (MAE) for each cell type was calculated as this measure has recently been established as a powerful predictor for the comparison of cell type deconvolution algorithms [31]. Overall, all four deconvolution algorithms performed comparably well on our test samples and were able to accurately discriminate cell types based on the selected ctDMPs. The CP method implemented in RnBeads performed best for the prediction of T-cell (MAE: 0.02) and monocyte (MAE: 0.03) fractions, whereas B cells were best predicted by EpiDISH (CBS; MAE: 0.04). For neutrophils EpiDISH (RPC), MeDeCom and RnBeads (CP) worked similarly well (MAE: 0.07; **Figure 2C**). The power to predict differences in cellular composition using the selected ctDMPs for cell type deconvolution was further highlighted by an in-depth analysis of the monocytic compartment: all four algorithms predicted an increase in the monocyte fraction in unsorted total bone marrow of JAK2V617F mutant mice (**Figure 2D**), which could be confirmed by flow cytometry (**Figure 2E, F**). Of note, the MEP fraction was systematically predicted to be higher than the fraction of MEPs as measured by FACS, resulting in a relatively high MAE (MAE: 0.22-0.27; **Figure 2C**). A likely explanation for this finding could be that MEP-specific methylation programs also identify other nucleated erythrocyte progenitor cells present in the bone marrow. It is well established that nucleated erythrocyte precursors make up ∼20% of all nucleated bone marrow cells which would fit well with the predicted values [32]. This interpretation is further supported by the observed increase in the predicted fraction of MEPs in both JAK2V617F knock-in bone marrow samples, which is in line with published data demonstrating an increase in erythrocyte progenitors in this model [33].

**Figure 2.**
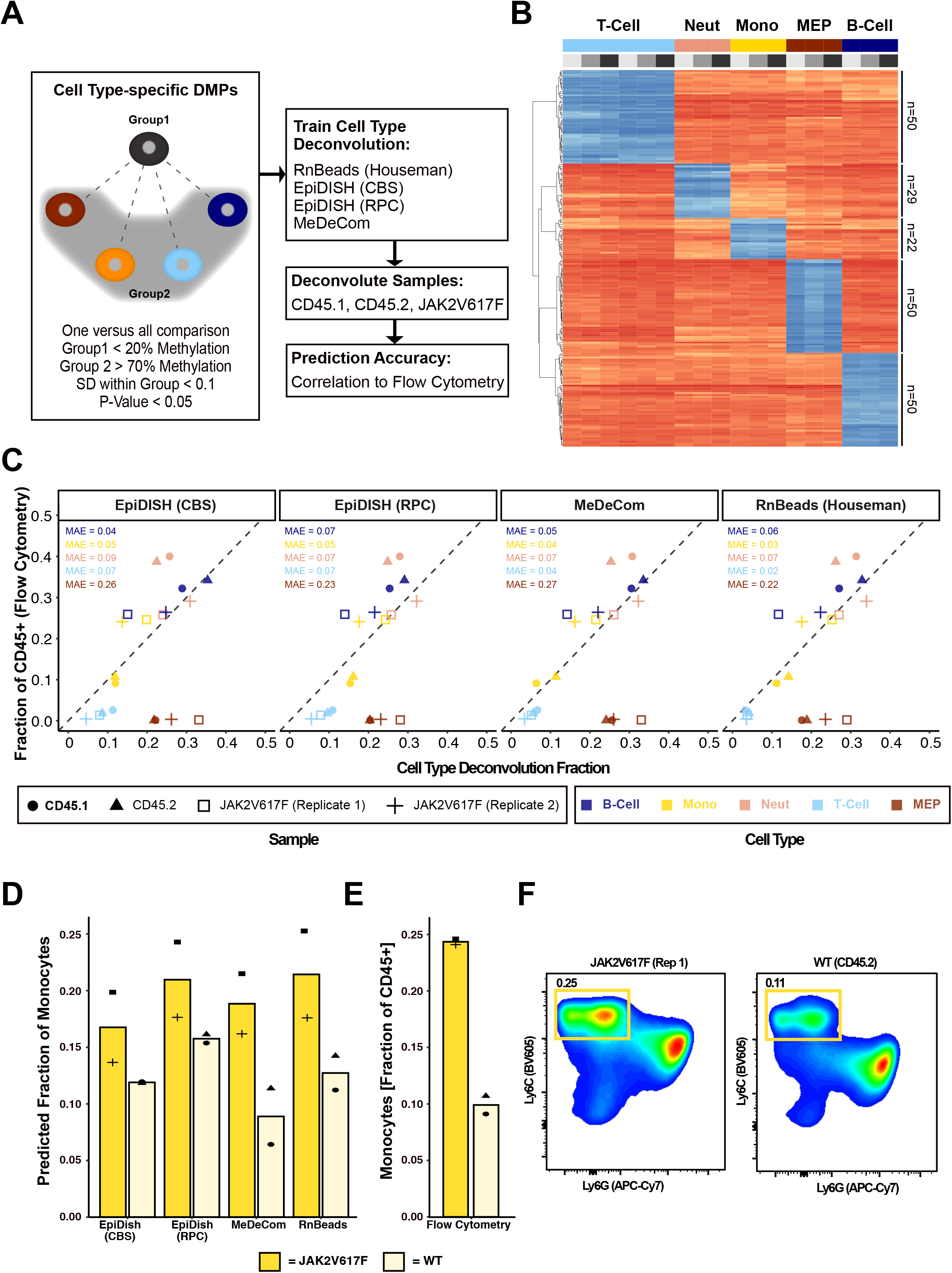
Cell type deconvolution. **(A)** Schematic overview of the cell-type deconvolution pipeline. Cell-type specific DMPs (ctDMPs) were determined in a one-versus-all manner using a stringent filtering strategy. The identified ctDMPs were used as input for 4 different cell type deconvolution algorithms. The predicted fractions for C57BL/6 wildtype (CD45.1 & CD45.2) and JAK2^V617F^-mutant mice (n=2) were compared to flow cytometry measurements to assess the prediction accuracy. **(B)** Heatmap depicting the DNA methylation β-values for the ctDMPs. **(C)** Scatter plot comparing the predicted and measured cell type fractions. The mean absolute error (MAE) was calculated for each cell type and algorithm. **(D)** Barplot showing the predicted monocyte fractions resulting from the different deconvolution algorithms for wildtype (CD45.1 & CD45.2) and JAK2^V617F^-mutant mice. **(E+F)** Flow cytometry data assessing the fraction of monocytes of all CD45+ cells. Data are shown as barplot **(E)** and density plot to visualize the flow cytometry gating **(F)**.

In conclusion, we established an enhanced strategy to define ctDMPs from MMBC array data which encompasses the most abundant hematopoietic cell types present in the bone marrow. These ctDMPs allow accurate cell type deconvolution independent of the chosen algorithm, and facilitate an orientating analysis of aberrant hematopoiesis from bulk bone marrow samples.

### DNA methylation dynamics during murine hematopoietic differentiation

Next, we aimed to comprehensively investigate DNA methylation programming during hematopoiesis. We calculated pairwise differential methylation between HSPCs and each of the remaining eight cell types analyzed to determine all CpGs that show dynamic methylation across hematopoietic differentiation (“differentiation-dynamic differentially methylated probes”; diffDMPs). In total, we identified 37,512 diffDMPs (**Supplementary Table 5; Supplementary Table 6**), of which 20,195 were found to be differentially methylated in more than one cell type (*“shared diffDMPs”*) while 17,317 diffDMPs were exclusively identified in a single cell type (*“unique diffDMPs”*). During hematopoietic differentiation, DNA methylation changes were predominantly characterized by a loss of DNA methylation with 88% (15,278) of the unique diffDMPs and 74% (14,985) of the shared diffDMPs showing hypomethylation compared to HSPCs (**Figure 3A**). Gain of DNA methylation during differentiation from HSPCs to differentiated cell types was almost exclusively observed in cells of the lymphoid lineage (**Figure 3A**). The vast majority of diffDMPs determined for CMPs (100%) and GMPs (99.93%) were shared diffDMPs. In contrast, unique diffDMPs were typically identified in MEPs and in differentiated cell types: in MEPs 49% of diffDMPs were unique to this cell type, followed by CD4+ T cells (33% unique diffDMPs), neutrophils (21% unique diffDMPs) and monocytes (11% unique diffDMPs).

**Figure 3.**
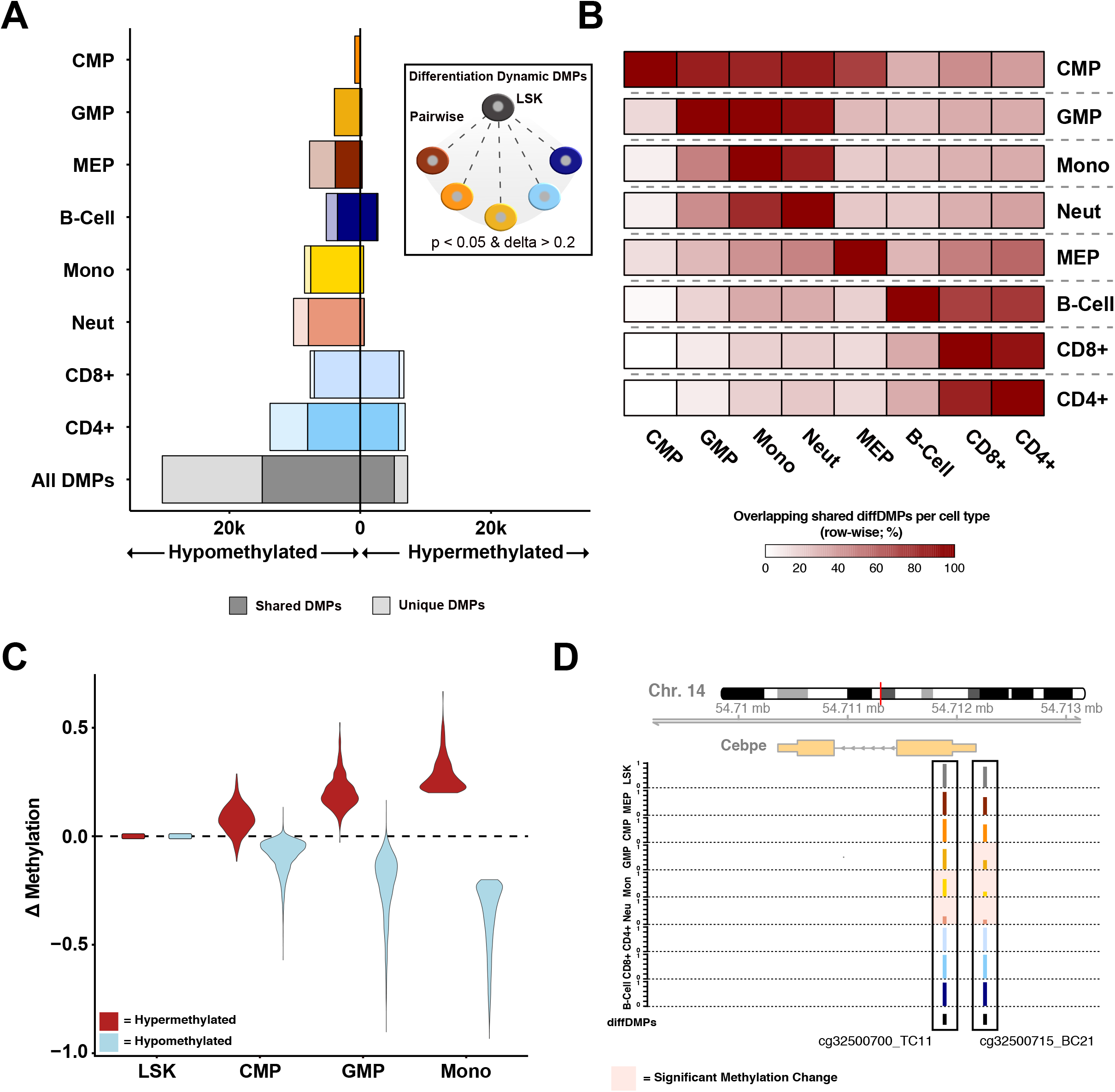
DNA methylation changes underlying murine hematopoietic differentiation. **(A)** Differentiation dynamic DMPs (diffDMPs; p<0.05 and delta methylation >0.2) were computed between LSK cells and all other cell types. Hypo- and hypermethylation refer to loss and gain of DNA methylation in the respective cell type as compared to LSK cells. The number of hypo- and hypermethylated probes per cell type is depicted as a bar plot. Each set of DMPs has been divided into shared DMPs (also differentially methylated in another cell type) and unique DMPs. **(B)** Heatmap showing the percentage of shared DMPs across cell types (row-wise). **(C)** Violin plot showing the DNA methylation change of monocyte-specific diffDMPs from LSKs to cell types of the myeloid lineage. The diffDMPs were stratified as hypo- or hypermethylated based on the DNA methylation difference to LSK cells. The DNA methylation differences (Δ methylation) were calculated by subtracting β-values of LSKs from each cell type. **(D)** Locus plot depicting the genomic region of the myeloid transcription factor *Cebpe*. Methylation β-values for the two diffDMPs identified in this region are shown as barplots and significant methylation changes are indicated with shaded background.

Next, we investigated whether shared diffDMPs identify DNA methylation programs that are “inherited” along hematopoietic differentiation trajectories. To disentangle the relationships of shared diffDMPs across the different cell types, we calculated the percentage of overlapping diffDMPs across cell types (**Figure 3B**, **Supplementary Table 7**). A large fraction of diffDMPs identified in CMPs overlapped with diffDMPs identified in other myeloid cell types and in MEPs, indicating that DNA methylation changes at these sites are faithfully propagated along the myeloid and the erythrocyte/megakaryocyte differentiation axis (**Figure 3B**, **Supplementary Figure 4, Supplementary Table 7**). A similar pattern was observed for diffDMPs identified in GMPs, where we found that the majority of diffDMPs were shared with monocytes (98%) and neutrophils (94%) whereas a much smaller proportion of GMP diffDMPs were shared with CMPs (18%). Of note, only very few diffDMPs identified in cells of the myeloid lineage were shared with lymphoid cell types and vice versa showing the divergence of these lineages at the epigenetic level. Interestingly, 75% of diffDMPs identified in CMPs were shared with MEPs, whereas only 29% of GMP diffDMPs were shared with MEPs (**Figure 3B**). This observation is in line with a previously described erythroid/megakaryocytic-priming of a CD55+ progenitor cells which constitute a substantial fraction of immunophenotypic CMPs but are virtually absent in GMPs [25].

These findings suggested that cell type-specific DNA methylation patterns are established in a progressive manner during differentiation and hence might serve as a molecular barcode to decipher differentiation trajectories *in vivo*. This can be exemplified by looking at the DNA methylation changes of monocyte-specific diffDMPs along the myeloid differentiation axis. Thus, diffDMPs which are hypomethylated in monocytes show the highest DNA methylation level in HSPCs and progressively lose methylation along the myeloid differentiation trajectory until they reach the lowest level in monocytes. Vice versa, diffDMPs that are hypermethylated in monocytes show the lowest DNA methylation level in HSPCs and continuously gain methylation along the monocytic differentiation axis (**Figure 3C**). The progressivity of the DNA methylation changes observed during hematopoietic differentiation appears to not only be a result of a progressive loss of DNA methylation at single CpG positions, but also involves the processive recruitment of neighboring CpGs (**Figure 3D**).

### diffDMPs reveal novel candidate *cis*-regulatory elements in the murine genome

The observed “inheritance” of DNA methylation patterns acquired during differentiation suggested a potential role of diffDMPs in the regulation of hematopoiesis. To confirm this hypothesis, we analyzed the enrichment of binding motifs from known hematopoietic transcription factors (TFs) in hypomethylated diffDMPs. We found enrichment of KLF TF-family motifs in MEPs and enrichment of GATA TF-family motifs in MEPs and CMPs (**Figure 4A**). Myeloid lineage-specification was accompanied by an enrichment of CEBP and PU.1 TF motifs in hypomethylated diffDMPs found in CMPs and GMPs. Terminally differentiated cell types showed characteristic enrichment patterns of TF binding motifs. Thus, diffDMPs hypomethylated in monocytes were enriched for IRF motifs; diffDMPs hypomethylated in CD4+ and CD8+ T cells were enriched for TCF and LEF1 motifs; and diffDMPs hypomethylated in B cells showed EBF and PAX motif enrichment. This enrichment of lineage-specific TF binding motifs in hypomethylated diffDMPs suggests that these sites might have *cis-* regulatory potential during hematopoietic differentiation. Hence, these sites could indicate the location of *cis-*regulatory elements (CREs) in the murine genome.

**Figure 4.**
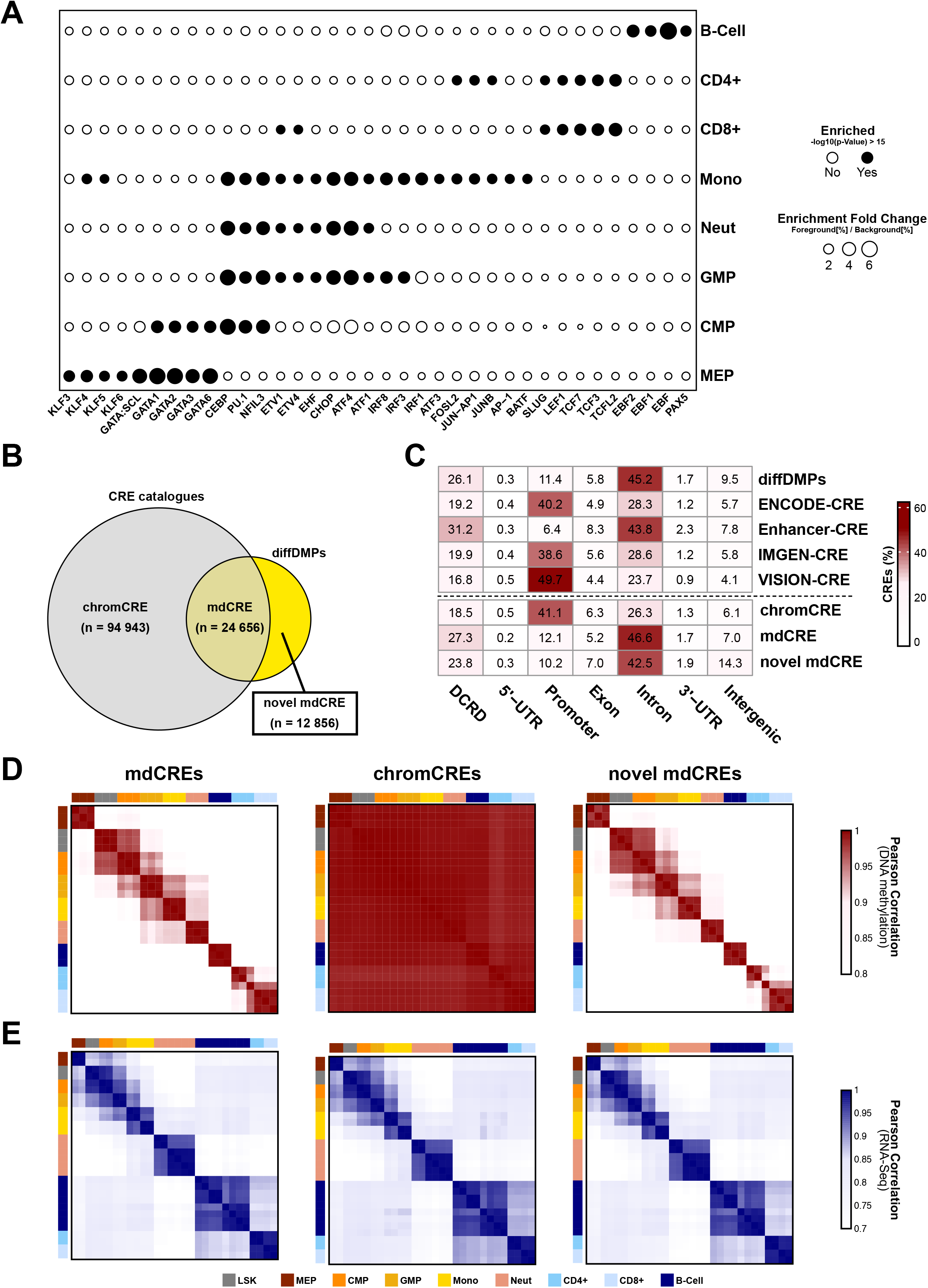
Differentiation dynamic DMPs mark candidate *cis-*regulatory elements. **(A)** Enrichment of transcription factor binding motifs in hypomethylated diffDMPs (±50 bp) identified in the different cell types. Enriched fold change (circle size) and significant enrichments (filled circle) are shown. **(B)** Venn diagram depicting the overlap of diffDMPs (n=37,512) with known CREs (n=119,599 MMBC CpGs overlapping with known CREs). chromCREs: chromatin-regulated CREs, i.e. CpG sites overlapping with known CREs but without significant methylation changes in hematopoietic differentiation; mdCREs: methylation-dynamic CREs, i.e. differentially methylated CpGs overlapping with known CREs; novel mdCREs: novel methylation-dynamic CpGs, i.e. differentially methylated CpGs <u>not</u> overlapping with known CREs. **(C)** Genomic localization for probes overlapping with the different CRE catalogues and subsets based on the distance to annotated genes. DCRD: distant *cis*-regulatory domain; 5’-UTR: 5’ untranslated region; 3’-UTR: 3’ untranslated region. **(C)** Pearson correlation heatmap of methylation β-values for probes of the different CRE subsets (mdCREs, chromCREs & novel mdCREs). **(E)** Pearson correlation heatmap based on RNA-seq data of genes associated with the different CRE subsets (mdCREs, chromCREs & novel mdCREs).

To further investigate this hypothesis, we compared the diffDMPs to four previously published CRE catalogues: candidate CREs from the ENCODE project (ENCODE-CRE; n=339,815)[18]; active hematopoietic enhancers (Enhancer-CRE; n=48,396)[34]; open chromatin regions (OCRs) identified in the Immunological Genome Project (IMGEN-CRE; n=512,595)[16]; and CREs from the validated systematic integration of hematopoietic epigenomes (VISION) project (VISION-CRE; n=205,019)[17]. A significant number of CpGs covered by the MMBC array overlapped with the individual CRE catalogues (min: 1.96 x 10^4^, max: 7.78 x 10^4^ CpGs), covering between 11% and up to 27% of the CREs defined by these catalogues (**Supplementary Figure 5A; Supplementary Table 8**). In total, 119,599 CpGs represented on the MMBC array overlapped with known CREs as defined by the aforementioned catalogues (**Figure 4B**). When considering all 37,512 diffDMPs, we found that 24,656 diffDMPs (66%) overlap with known CREs and hence were defined as methylation-dynamic CREs (mdCREs; **Figure 4B; Supplementary Figure 5B**). Importantly, the non-overlapping fraction of 12,856 diffDMPs (34%) constituted the third-largest region set found in a comprehensive overlap analysis of the individual CRE datasets (**Supplementary Figure 5B**). This suggests that this subset of diffCpGs might indicate the presence of novel, yet unrecognized, hematopoietic CREs and therefore were named “novel methylation-dynamic CREs” (novel mdCREs; **Figure 4B**; **Supplementary Figure 5B; Supplementary Table 6**). The remaining 94,943 CREs (79%), which did not overlap with diffDMPs, were termed chromatin-dynamic CREs (chromCREs). These chromCREs might represent regions which are either not regulated at all during hematopoiesis or they are dynamically regulated exclusively at the chromatin level and do not show significant DNA methylation changes.

To investigate potential differences in genomic features captured by the different catalogues, we analyzed the genomic localization of diffDMPs and of the different CRE subsets relative to neighboring genes. We found that chromCREs as well as ENCODE-, IMGEN- and VISION-CREs are enriched in promoter regions (38%-62%; **Figure 4C**). In contrast, diffDMPs, Enhancers, mdCREs and novel mdCREs are enriched in intronic regions (43%-47%) and distant *cis-* regulatory domains (DCRDs; 24%-31%; **Figure 4C**). The observed similarity in genomic distribution of diffDMPs and known hematopoietic enhancers suggested that diffDMPs, as a whole might identify both known (mdCREs) as well as novel (novel mdCREs) hematopoietic enhancers.

We next computed a correlation matrix of DNA methylation between all cell types to analyze how DNA methylation at these CRE sets could preserve cell type and lineage identity. This analysis revealed high correlation across closely related cell types and within the same lineages for mdCREs and novel mdCREs, whereas for chromCREs the observed DNA methylation patterns hardly allowed discrimination of cell types and lineages (**Figure 4D, Supplementary Figure 6A&B**). This analysis confirmed that chromCREs, in contrast to mdCREs and novel mdCREs, are not regulated dynamically at the level of DNA methylation during hematopoietic differentiation. To get an estimate of the functional activity of these CRE subsets in hematopoiesis, we assessed the expression of associated genes in publicly available RNA-seq datasets from murine hematopoietic cells. We annotated all mdCREs, chromCREs and novel mdCREs to the closest gene and correlated the expression of all associated genes across different cell types. This analysis revealed high correlation of gene expression across related cell types independently of whether they had been associated with mdCREs, chromCREs or novel mdCREs (**Figure 4E**). This indicated that all CRE subsets identify relevant *cis*-regulatory elements, but further suggested that the different CRE subsets might be subject to different mechanisms of epigenetic regulation. For instance, chromCREs do not show prominent changes in DNA methylation and therefore might be regulated mainly at the chromatin level. The mdCREs show dynamic regulation at the level of DNA methylation and, presumably, also at the chromatin level, since they had been identified using chromatin-based methods. In contrast, novel mdCREs are exclusively identified in the present study based on dynamic DNA methylation changes during hematopoietic differentiation while previous studies investigating chromatin changes in the hematopoietic system failed to identify these regulatory elements. In summary, our analysis revealed three different classes of CREs: i) CREs which have been identified by chromatin-level characteristics but lack DNA methylation dynamics (chromCREs); ii) CREs which have been identified by chromatin-level characteristics and which show dynamic DNA methylation changes during hematopoietic differentiation (mdCREs); and iii) novel CREs which are so far exclusively characterized by dynamic DNA methylation (novel mdCREs). Importantly, the 12,856 novel mdCREs have not been described in previous datasets, which underlines the importance of DNA methylation analysis for the functional annotation of genomes. This finding further suggests that dynamic regulation of DNA methylation might play a key role as an epigenetic regulatory layer in a subset of *cis*-regulatory elements.

### Novel mdCREs exhibit cell type-specific DNA methylation programs that can be annotated to putative target genes

Having demonstrated on a global scale that DNA methylation of the novel mdCREs correlates with hematopoietic cell-type identity, we performed unsupervised hierarchical clustering to characterize the DNA methylation programs encoded by the 12,856 novel mdCREs in more detail. This clustering identified nine lineage-specific DNA methylation programs which allowed the discrimination of all the cell types analyzed (**Figure 5A&B**; **Supplementary Figure 7 A&B; Supplementary Table 6**). Remarkably, eight clusters showed loss of DNA methylation from LSK to more differentiated cell types, whereas one cluster (cluster 7) predominantly showed gain of DNA methylation in differentiated lymphoid cells, with a maximum reached in CD4+ and CD8+ T cells (**Figure 5A&B**; **Supplementary Figure 7 A&B**). Specifically, MEPs exhibited loss of DNA methylation in cluster 4; CD4+ T cells demonstrated loss of DNA methylation in clusters 1 and 2; both CD4+ and CD8+ T cells were characterized by DNA methylation loss in cluster 9; while the entire lymphoid lineage displayed loss of DNA methylation in cluster 3. Cluster 8 specifically showed loss of DNA methylation in B cells, and cells from the myeloid lineage exhibited loss of DNA methylation in clusters 5 and 6. This coordinated methylation programming of CpG sites observed in the novel mdCREs further supported the idea that these CpG sites identify regions which exert important regulatory functions during hematopoietic differentiation.

**Figure 5.**
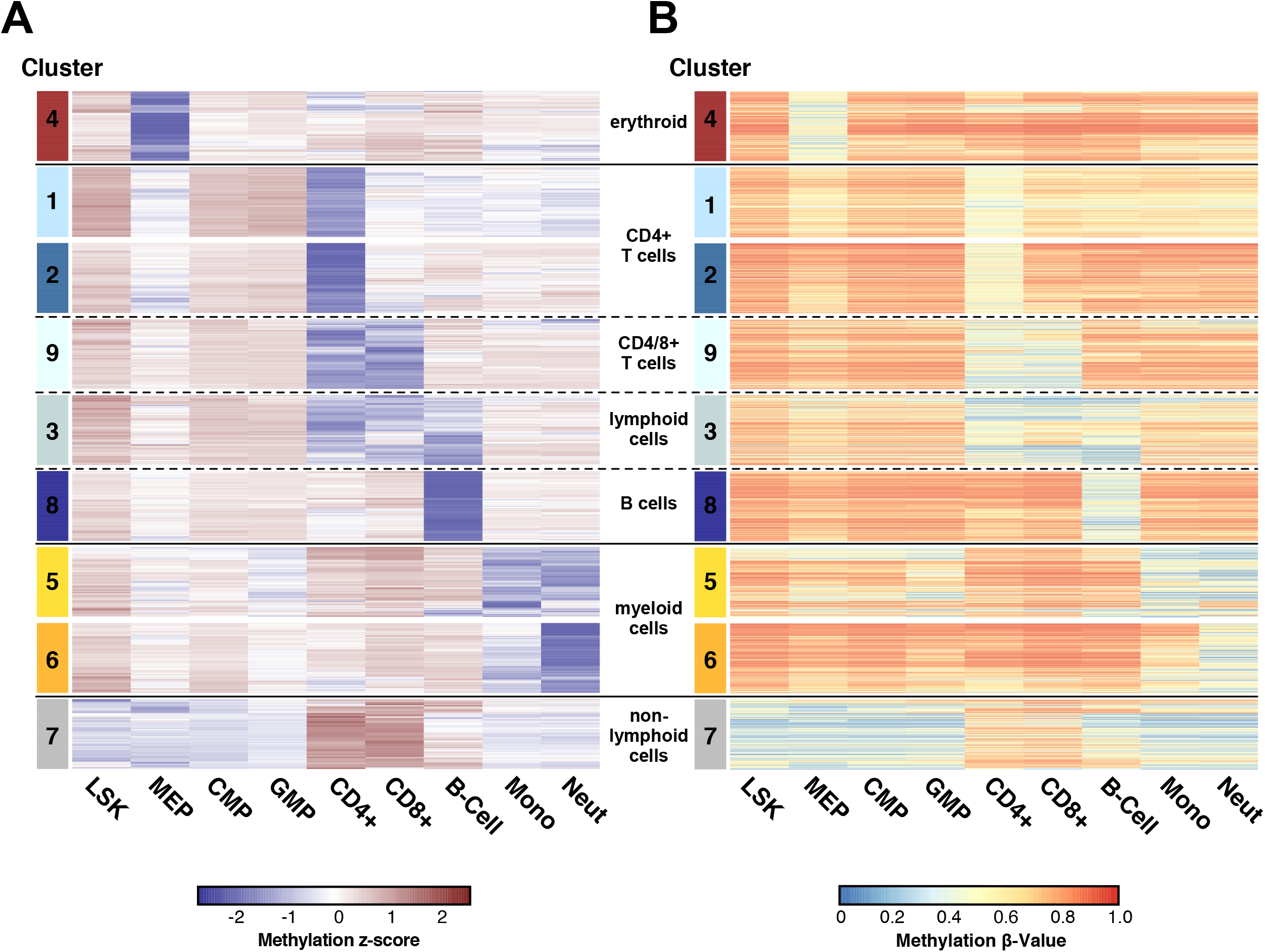
Cell type-specific DNA methylation programs of novel mdCREs. DNA methylation β-values of novel mdCREs were z-score transformed and hierarchically clustered using Euclidean distance and Ward’s method. This strategy identified 9 different clusters with cell type specific DNA methylation patterns. For display purposes 350 CpG sites were randomly selected from each cluster. The z-score transformed methylation β-values of these sites were again clustered within each cluster. Plotted are the z-score transformed data **(A)** as well as the corresponding absolute β-values **(B)** of the same CpG sites in the same order. Please refer to Supplementary Figure 7 for heatmaps representing all novel mdCRE CpGs.

Next, we aimed to identify genes that are likely regulated by the novel mdCREs. Since recent reports indicated that the regulatory potential of CREs may expand to a megabase scale [35], we developed an heuristic approach to infer putative functional CRE-gene pairs based on a systematic integrative analysis of DNA methylation and gene expression patterns (**Figure 6A**). First, all transcription start sites (TSSs) within 1Mb distance of the 12,856 novel mdCREs were determined and selected as “candidate associations”. A total of 406,002 TSSs were identified as candidate association, meaning that a median of 25 TSSs (min=1; max=167) were assigned to each novel mdCRE. For each candidate association, a linear model was trained assuming that the DNA methylation β-values predict the gene expression levels in the same cell populations, resulting in the identification of 1,445 significant candidate associations (Benjamini-Hochberg adjusted correlation test p-value <0.01). As an additional filter, we used the slope of the linear model as a surrogate for the effect size of the regulation. Using these stringent criteria, we found 843 significant novel mdCRE-gene pairs, which we propose as strong candidates for further functional validation studies (**Supplementary Table 9**). Looking at the genomic distribution of these 843 novel mdCRE-gene pairs, we observed that only 39 (4.6%) novel mdCREs were located within 5kb to the TSS, whereas the majority of novel mdCREs were found to be evenly distributed within 1Mb distance up- or downstream of the TSS (**Figure 6B**) and were associated with a single gene (**Figure 6C**). Five of the novel mdCREs were each associated with more than 20 genes. For example, the methylation of one CpG site correlated with the expression of the T-cell receptor beta chain gene cluster, indicating that one mdCRE has a regulatory potential for several genes with similar functions. The *Thy1* locus, in contrast, is an example for a gene locus which is associated with many novel mdCREs. In total, 35 novel mdCRE-gene associations were identified within a 1Mb distance of the *Thy1* TSS, of which 10 associations fulfilled the correlation and effect size criteria (**Figure 6D**). To further investigate whether these associated novel mdCREs could function as distal regulatory elements, we analyzed HiC data of CD8+ T cells and found that the associated novel mdCREs overlapped with high contact domains suggesting a possible physical interaction between these novel mdCREs and the *Thy1* TSS (**Figure 6E**).

**Figure 6.**
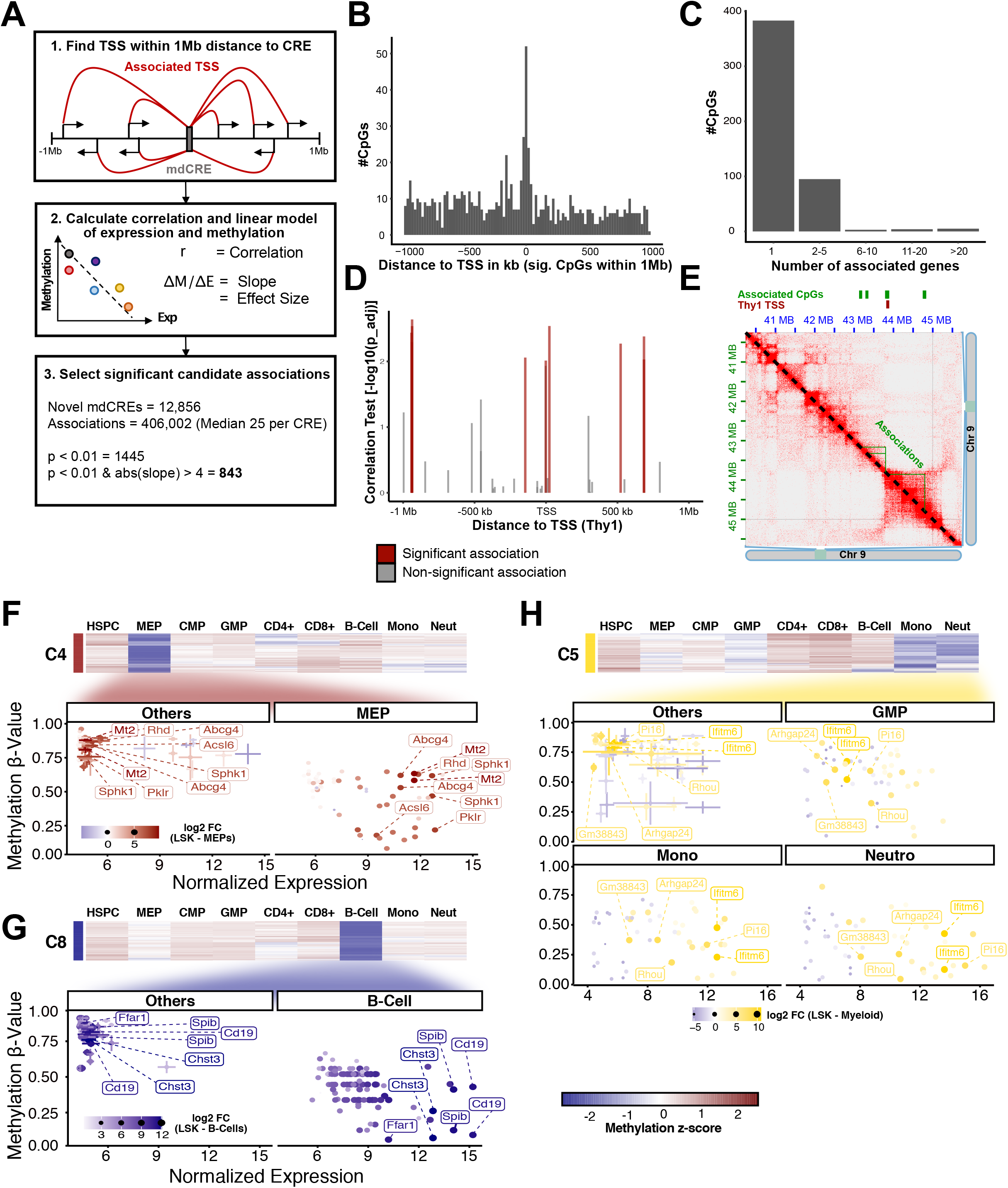
Hypomethylation of novel mdCREs is associated with lineage specific gene expression patterns. **(A)** Strategy to annotate novel mdCREs to putative target genes. All transcriptional start sites (TSS) within an 1Mb radius to novel mdCREs were mapped. Next, Pearson correlation and a linear model (surrogate of DNA hypomethylation effect size) between gene expression of associated genes and methylation β-values of novel mdCREs was calculated. All associations with an FDR-adjusted correlation p-value <0.01 and an absolute slope of >4 were selected as significant candidate associations. **(B)** Distribution of significant candidate associations around TSS. **(C)** Histogram showing the number of associated genes per novel mdCRE. **(D)** Barplot of FDR-adjusted correlation p-values for novel mdCREs within a 1Mb distance to the *Thy1* TSS and the respective gene expression. Significant candidate associations are colored as red bars. **(E)** HiC contact map for the *Thy1* locus showing the localization of significant associated novel mdCREs and the *Thy1* TSS. **(F-H)** Scatter plots show the methylation β-values and gene expression values (vst-transformed counts) for associated novel mdCRE-gene pairs within the different novel mdCRE methylation clusters. Methylation z-score heatmaps for the CpG sites in the respective clusters are shown above the scatter plots. Cell populations with cluster-specific hypomethylation (**F,** MEPs for cluster 4; **G,** B-cells for cluster 8; **H,** myeloid lineage for cluster 5) have been compared to all other cell types (gene expression and methylation β-value range in other cell types depicted by error bars). The coloring and point size show the log2 fold-change of a differential expression test between the respective cell type with the strongest hypomethylation and LSK cells.

Next, we performed a systematic analysis of the cell type- and lineage-specific gene expression changes and their association with DNA methylation changes of the novel mdCREs. To do so, the expression of genes which are annotated as candidate associations within the lineage-specific clusters of co-regulated CpG sites was analyzed (**Figure 6F-H**; **Supplementary Figure 8**). Genes associated with the erythroid cluster 4 revealed low methylation and high gene expression levels in MEPs. In contrast, all other populations showed high methylation and low or decreasing gene expression values (**Figure 6F**). Known erythroid marker genes like *Pklr* or *Sphk1* were among the cluster 4 associated genes which showed the strongest increase in gene expression accompanied by loss of DNA methylation in MEPs. Similar results were obtained for the B cell specific cluster 8 where B cell expression dynamics was compared to all other cell types (**Figure 6G**). In this cluster, B cell marker genes such as *Cd19* or *Spib* demonstrated loss of DNA methylation which was paralleled by an induction of gene expression. In turn, all other cell types showed stable DNA methylation patterns, accompanied by low gene expression levels. A more complex situation was observed for genes associated with myeloid cluster 5 (**Figure 6H**). In this cluster, the expression of myeloid (monocyte + neutrophil) marker genes was analyzed for GMPs, neutrophils and monocytes, as these cell types showed cluster-specific hypomethylation. Compared to all other cell types, myeloid marker genes showed initiation of DNA methylation loss in GMPs whereas the strongest hypomethylation was observed in terminally differentiated myeloid cells. This loss of DNA methylation was accompanied by a strong increase in expression of these genes from GMPs to monocytes/neutrophils. Among those genes, *Ifitm6* showed a continuous induction from GMPs to neutrophils which is in line with the high expression of type I interferon response genes during neutrophil specification [36]. In summary, we developed an algorithm which allowed us to propose putative functional gene annotation for a subset of the novel mdCREs. In addition, we identified DNA methylation programs that are defined by co-regulated novel mdCREs and which are associated with hematopoietic-specific gene expression patterns that are progressively established during hematopoietic differentiation.

## DISCUSSION

Many biological insights into the distinct role of DNA methylation have emerged with the advent of array-based DNA methylation profiling technologies. These arrays offer an affordable, easy-to-use and robust platform for DNA methylation profiling of hundreds to thousands of samples. However, array-based technologies have not been available until recently for the study of murine methylomes. This has compromised many mechanistic and functional studies investigating the role of DNA methylation changes in mouse models.

The novel murine methylation bead chip (MMBC) array filled and allows to profile DNA methylation of 285k CpG sites across the genome. The potential of this method has recently been demonstrated by a mouse DNA methylation atlas encompassing MMBC data for distinct tissues, mouse strains, age groups and pathologies [37]. This study also demonstrated the robustness and high reproducibility of the DNA methylation measurements generated using MMBC. Hence, the MMBC array will help to accelerate the research of epigenetic plasticity in homeostasis and disease.

To establish a computational workflow for the analysis of MMBC datasets, we generated a comprehensive annotation of the MMBC array. This includes mapping of probes to nearest genes and an annotation of functional genomic elements such as promoters or distal cis-regulatory domains (DCRDs). Additionally, we expanded the commonly used and highly cited RnBeads framework by methods that allow the processing of MMBC data [22, 23]. This includes user-friendly functions for quality control, normalization, and differential methylation calling. Moreover, RnBeads generates automated reports which document analysis parameters and will thus enhance the reproducibility of MMBC data analyses.

In the present study, we chose to profile DNA methylation changes during murine hematopoiesis using the MMBC array. The advantage of this system is the well-defined differentiation landscape including the opportunity to isolate homogeneous cell populations using previously established cell surface markers. This allowed us to assess the robustness of the MMBC array by investigating this complex *in-vivo* differentiation system with a focus on the myeloid differentiation trajectory. We observed an accurate clustering of cell types based on global DNA methylation changes which indicated that the MMBC array is capable of determining cell type-specific DNA methylation programs across closely related cell types. Hence, we challenged the MMBC array to test whether it is a suitable tool for DNA methylation-based cell type deconvolution. Therefore, we have elaborated a strategy to determine highly cell type-specific DNA methylation signatures (ctDMPs) which were defined by the highest DNA methylation difference between cell types with a low intra-cell type variance at the same time. We hypothesized that these sites possess the potential to discriminate true epigenetic plasticity from an impure DNA methylation signal due to a heterogenous cell population. To test this, we generated DNA methylation data from mixed cellular populations and applied four different published cell type deconvolution algorithms. The predicted cell type fractions from each of the algorithms were comparable to the direct measurement of the cell type composition by flow cytometry. Moreover, all four algorithms were able to confirm the disease-specific imbalance in the cellular composition of the bone marrow of JAK2^V617F^-mutant mice. In summary, highly cell type-specific DNA methylation signatures can be identified with the MMBC array which allow an accurate estimation of cell fractions in healthy and diseased tissues, independent of the underlying deconvolution algorithm.

Besides the homeostatic state of a cell, the DNA methylation signature harbors information about its lineage history and potentially also insights into future fate decisions [38]. This implies that cell types of a certain lineage share DNA methylation signatures. To identify such programs, we considered the HSPC compartment as a common starting point in hematopoiesis and determined differentiation dynamic CpG sites (diffDMPs) for all cell types relative to HSPCs. We could identify three properties of these sites: i) diffDMPs of progenitor cells were shared with their downstream progeny; ii) the number of diffDMPs increased with cellular differentiation states; iii) the DNA methylation level of dynamic CpG sites progressively evolved towards differentiated cells. Taken together, this indicated that DNA methylation programs progressively and unidirectionally develop along hematopoietic differentiation trajectories. Importantly, since CpG sites can only exist in two DNA methylation states (methylated or unmethylated), intermediate DNA methylation ß-values have to arise from heterogeneity of the investigated sample. Consequently, progressive changes of DNA methylation ß-values within diffDMPs either indicate that the isolated cell populations are mixtures of cells with heterogenous differentiation states or that at the level of individual cells cellular commitment does not occur in a fully coordinated manner. However, this problem cannot be addressed with the currently available technologies as single-cell long read DNA methylation data would be required to answer this question.

Independent of the mechanisms underlying epigenetic heterogeneity at the population level, our study demonstrates pronounced remodeling of the DNA methylation landscape during hematopoiesis. These dynamics point towards a regulatory role of DNA methylation in the course of cellular differentiation. In fact, a substantial fraction of diffDMPs overlapped with previously described CREs which further underlines their regulatory potential. In addition to these methylation dynamic CREs (mdCREs), we identified 12,856 diffDMPs which have not been described as regulatory regions before. These regions represent novel candidates of methylation dynamic CREs (novel mdCREs). Of note, these single CpG sites do not represent the whole region of a CRE in the genome but rather flag genomic loci with regulatory potential. Importantly, we determined these sites by comparing them to comprehensive CRE catalogues which either included histone marks, open chromatin sites, or a combination of both [16–18, 34]. However, none of these catalogues takes DNA methylation as a regulatory mark into account. We demonstrated that mdCREs can be clustered into programs with cell type-specific DNA methylation patterns and that their methylation status strongly correlates with expression programs of neighboring genes. This confirms that we generated a unique resource of novel DNA methylation-based regulatory elements which are likely involved in the regulation of murine hematopoiesis and which are not restricted to CpG-dense regions as was the case for previous studies [9].

The functional annotation of CREs to their target genes is a central task in computational epigenomics. Previous studies have shown that interactions between CREs and genes can occur on a megabase scale [35]. In consequence, algorithms which focus solely on a distance based CRE-to-gene annotation will only include a subset of putative regulatory interactions.

To address this issue, we developed a data-driven annotation strategy which incorporates the DNA methylation status of CREs and the expression of genes in a region +/- 1 Mb of a TSS. We could observe regulatory candidate associations both in the close proximity of TSSs and in distant regions. The algorithm can in principle be expanded using different data sources like histone marks and will allow the functional annotation of CREs in complex biological systems. While the inferred candidate associations propose a biological function of CREs and seem to be biologically meaningful, further studies e.g. using massive parallel reporter assays (MPRA) will be required to confirm these interactions.

In summary, we generated a reference atlas of dynamic DNA methylation changes during murine hematopoietic differentiation using the recently released MMBC array. This atlas includes a comprehensive list of CpG sites with dynamic DNA methylation during murine hematopoiesis and candidate associations for novel regulatory elements. Moreover, we developed a computational pipeline for a fast, robust, reproducible and user-friendly analysis of MMBC data and propose and analysis workflow which can be applied to various tissues and disease models and thus constitutes a major resource for epigenetic studies in the murine system.

## METHODS

### Mice and ethics statement

C57BL/6J (Cd45.2) and B6.SJL-Ptprc^a^ Pepc^b^/BoyCrl (Cd45.1) mice were bred in-house at the German Cancer Research Center (DKFZ) under specific pathogen-free (SPF) conditions. Mice at the age of six to sixteen weeks were used for all experiments.

In addition, bone marrow samples from mice expressing the conditional JAK2^V617F^ mutant allele were used [33]. The mutant allele was induced by a Vav1-Cre recombinase. Bone marrow was harvested from JAK2V617F-expressing animals (JAK2^VF/+^ Vav-Cre+) and the respective JAK2-wildtype (JAK2^+/+^ Vav1-Cre+) littermate controls at 10-12 weeks of age. All mouse experiments were approved by local authorities according to German and European guidelines.

### Bone marrow isolation and cell sorting

Tibiae, femora, iliae, vertebrae, sterna and humeri of sacrificed mice were isolated and crushed three times in Iscove’s Modified Dulbecco’s Medium (IMDM, Gibco) using a mortar and pestle. The supernatant was filtered through 40 µm cell strainers (Falcon) and bone marrow cells were pelleted by centrifugation. Red blood cell lysis was performed by adding ACK lysis buffer (Lonza). The bone marrow cellularity was determined by counting with a veterinary hematology analyzer (scil Vet abc Plus+, scil). Lineage depletion was performed for the isolation of hematopoietic stem and progenitor cells (LSK, CMP, GMP and MEP) and T cells (CD4 T cells and CD8 T Cells). In brief, biotinylated lineage antibodies were added to the isolated bone marrow cells (HSCPs: CD5, CD8, B220, CD11b, Gr1 and Ter-119; T cells: CD11b, CD16/32, B220 and Ter-119; manufacturer details in **Supplementary Table 2**) and the labeled cells subsequently incubated with Mouse Depletion Dynabeads (Invitrogen). Purification of lineage-negative cells was performed using a DynaMag-15 magnet (Invitrogen). Cells were stained according to previously established gating strategies for the murine hematopoietic system [39, 40](**Supplementary Figure 1; Supplementary Table 2**). Cell sorting was performed using a BD FACS Aria II or III cell sorter (BD Biosciences). In total 3 x 10^5^ cells were sorted per tube and snap-frozen on dry ice.

### Flow Cytometry

For flow cytometry, isolated bone marrow cells were stained with specific panels for the identification of hematopoietic cell lineages (**Supplementary Table 2; Supplementary Figure 1**). All measurements were performed on a LSRII flow cytometer (BD Biosciences).

### DNA isolation and Infinium mouse methylation bead chips (MMBC)

Snap-frozen cell pellets were thawed on ice and DNA was isolated using a QIAamp DNA Micro Kit (QIAGEN) according to the manufacturer’s instructions. DNA concentrations were measured using the Qubit dsDNA HS Assay Kit (Invitrogen). Integrity of genomic DNA was verified by the DKFZ Genomics and Proteomics Core Facility and 100-250 ng subjected to DNA methylation analysis using Infinium Mouse Methylation BeadChip arrays (Illumina, San Diego, CA, USA).

### RnBeads-compatible annotation of the MMBC array

The *RnBeadsAnnotationCreator* package (https://github.com/epigen/RnBeadsAnnotationCreator) was modified to generate an RnBeads-compatible annotation of the MMBC array for the mm10 reference genome. In short, the *Infinium Mouse Methylation Manifest File* (version 1.0) was downloaded and divided into assay and control probes. The assay probes included 287,050 probes of which 284,860 probes were annotated to bind in a CG-context, 838 in a CH-context and 1,352 in a SNP-context. Among those, 24,860 probes (22,670 CG, 838 CH and 1,352 SNP) were marked as “MFG_Change_Flagged” indicating manufacturing related performance problems. These probes with additional 26 probes binding on the mitochondrial genome (chrMT) were excluded from the reference annotation. This resulted in 262,164 probes which were considered for the RnBeads DNA methylation analysis workflow. The control probes were annotated as bisulfite-conversion, specificity, non-polymorphic or negative control probes, with either an expected high- or background-intensity as provided by Illumina. The resulting probe annotations together with further mm10 genome annotations which are required for the RnBeads workflow are available as *RnBeads.mm10* package from Bioconductor [24].

### DNA methylation array data processing

A pipeline for the analysis of MMBC data was developed within the *RnBeads* Bioconductor environment (Release 3.1.4)[22–24]. In short, IDAT files were imported in *RnBeads* and a quality control report was generated to inspect data quality. Background subtraction (“rnb.bgcorr.subtr”) and subsequent intra-array dye bias normalization (“rnb.norm.scaling”) with an internal reference were performed. Unreliable probes (Greedycut algorithm with detection p-value < 0.01) and probes mapping to sex chromosomes were removed from the dataset. For unsupervised inspection of the data, a principal component analysis (PCA) was computed based on the 5000 most variable CpGs (mvCpGs) as determined by standard deviation.

### Functional annotation of the MMBC array

Gene and genomic region annotation were performed with *gtfanno 0.2.0* (https://github.com/stephenkraemer/gtfanno). For all probes, residence in the following genomic regions was considered (in order of precedence): i) promoter (1500 bp upstream to 500 bp downstream ofTSS); ii) 5’UTR or3’UTR; iii) intron or exon iv) distant *cis*-regulatory domain (DCRD; region +/-100 kb from TSS) v) intergenic (if no other category was met). Based on a recent large-scale study of the mouse immune system and its differentiation cascades, we used a window of +/-100kb around the TSS for annotating putative enhancer relationships [16]. Annotations were performed against Gencode (release M25 for GRCm38/mm10), considering only the principal isoform of protein coding genes (based on the APPRIS isoform annotations provided with Gencode M25)[41]. If a probe was annotated to several region classes, only annotations for the region class with the highest precedence were considered. Within the highest ranked region class for each probe, all possible gene annotations were kept. For example, a probe may reside in the promoter region of two genes. In this case, both gene annotations were kept. The distance of the probe to the TSS (for promoter and DCRD annotated probes) or to the center of the exon, intron or UTR is detailed in the annotation table to allow for further ranking in cases were multiple genes were annotated to a single probe (**Supplementary Table 1**). A documentation of the annotation workflow is publicly available (https://github.com/stephenkraemer/bead_chip_array_annotations).

### Comparison to data from hematopoietic cell types obtained by reduced representation bisulfite sequencing (RRBS) and tagmentation-based whole genome bisulfite-sequencing (TWGBS)

Tagmentation-based whole-genome bisulfite sequencing (TWGBS) data from LSK subpopulations (GSE146907) and reduced representation bisulfite sequencing data from hematopoietic progenitor and differentiated cell types (RRBS; https://medical-epigenomics.org/papers/broad_mirror/invivomethylation/index.html) from two previous publications was downloaded (**Supplementary Table 3**)[9, 42]. TWGBS data were processed using the *Methrix* Bioconductor package [43]. All autosomal CpG sites overlapping with the MMBC array with a minimal coverage of 10 reads in at least two replicates were considered for further analysis. For analysis of the RRBS data, a liftover from mm9 to mm10 was performed using the “liftOver” function from the *rtracklayer* R package and CpG sites overlapping with the MMBC array were determined [44]. For RRBS correlation analysis, all autosomal CpG sites with a coverage >20 reads were considered. Replicates from the same cell type were summarized by mean and plotted against MMBC data using the *ggpointdensity* package (https://github.com/LKremer/ggpointdensity).

### Unsupervised phylogenetic analysis

An unsupervised DNA methylation based phylogenetic reconstruction of the murine hematopoietic hierarchy was performed by extracting the 5000 mvCpGs from the dataset and calculating the Manhattan distance. The “fastme.bal” function from the *ape* R package was applied to infer phylogenetic trees based on a minimal evolution algorithm and trees were plotted using the ape “plot.phylo” function [45].

### Cell type deconvolution

To determine a set of CpG sites that could predict the composition of differentiated hematopoietic cell types in a tissue with a high accuracy, we calculated differentially methylated CpG sites in a one-versus-all fashion between T cells, B cells, Monocytes, Neutrophils and MEPs. CpG sites with a maximum methylation of 20% in the query cell type and a minimum methylation of 70% in all other cell types with a standard-deviation per group <10% and a false-discovery rate adjusted p-value < 0.05 were considered as cell type-specific DMPs (ctDMPs). These ctDMPs were then sorted based on the mean methylation difference (Δmeth) between the two groups and the top 50 CpG sites with the highest Δmeth were used for cell type decomposition (**Supplementary Table 4**). The cell type contribution in the bone marrow of CD45.1, CD45.1 and JAK2^V617F^-mutant mice was predicted using different established algorithms: Houseman’s constrained projection method (*RnBeads* implementation), *EpiDISH* robust partial correlations (RPC) and Cibersort (CBS) mode as well as the reference-free non-negative matrix factorization (NMF)-based method *MeDeCom* [27–30]. For *MeDeCom*, a parameter search was conducted to find the best LMCs (2-10) and λ-Value (Settings: NINIT=10, NFOLDS=10, ITERMAX=100, NCORES=20). The best results based on evaluation of the cross-validation error were achieved for 5 LMCs and a λ-Value of 0.01. The fractional cell-type contribution was calculated for the different samples and compared to flow-cytometry data obtained from the same samples. The mean absolute error (MAE) was calculated using the *Metrics* R package (https://github.com/mfrasco/Metrics) and used as a measure of the predictive power for the different algorithms.

### Differential methylation calling

Differentially methylated probes (DMPs) between LSK cells and all other cell types were determined in a pair-wise manner using the *RnBeads* “rnb.execute.computeDiffMeth” function. CpG sites with a mean methylation difference of >20% and a false-discovery rate adjusted p-value < 0.05 were considered as differentiation dynamic DMPs (diffDMPs) for each cell type (**Supplementary Table 5**). The localization diffDMPs around the TSS of common lineage marker genes was plotted using the *Gviz* Bioconductor package [46].

### Enrichment of transcription factor binding motifs

Enrichment of known transcription factor binding motifs within each hypomethylated diffDMP set was calculated using the *HOMER* software (v4.8)[47]. In short, the *HOMER* “findMotifsGenome” function was used with each cell-type diffDMP set as input data and all remaining CpG sites from the MMBC array (after quality control filtering) as a background set. The size of the tested region was fixed as 50 bp upstream and downstream of the input locus. Enrichment was calculated as the percentage of target regions with a known motif divided by the percentage of background regions with the same motif.

### Overlap with known *cis*-regulatory elements

Four previously published CRE catalogues were analyzed for overlap with the MMBC array and with diffDMPs: ENCODE candidate CREs (ENCODE-CRE), VISION project hematopoietic CREs (VISION-CRE), hematopoietic enhancer (Enhancer-CRE) and open chromatin regions (OCRs) of the Immunological Genome (IMGEN) consortium (IMGEN-CRE)[16–18, 34]. All CREs mapping to autosomes were considered for the analysis. IMGEN-CREs were expanded by 125bp upstream and downstream as these sites were reported as 250bp width centered on the summit [16]. Enhancer-CREs intervals were extracted from **Supplementary Table 1** provided in Lara-Astiaso *et al.* 2014 [34]. The mm9 interval coordinates were translated to mm10 coordinates with the UCSC *liftOver* command-line program. We used the mm9ToMm10.over.chain.gz chain file provided in the UCSC genome browser database (https://hgdownload-test.gi.ucsc.edu/goldenPath/mm9/liftOver/mm9ToMm10.over.chain.g z).

The number of overlapping CpGs was plotted using the *UpSetR* package (https://github.com/cran/UpSetR). We defined three sets of candidate CREs for further analysis based on the overlap between the catalogues used: *“methylation dynamic CREs”* (overlap of diffDMPs with any CRE of the previous catalogues), *“chromatin CREs”* (regions identified in previous CRE catalogues which did not show DNA methylation dynamics in our MMBC dataset) and novel *“novel methylation dynamic CREs”* (novel mdCREs; diffDMPs which did not overlap with any of the other sets). The Pearson correlation of the methylation β-values for these CRE sets was calculated for all profiled cell types.

### Clustering of candidate CREs

The mean methylation β-values of all novel mdCREs were calculated across all biological replicates of each cell type and were then transformed to z-scores. For unsupervised hierarchical clustering we used Ward’s method on the squared Euclidean distance. We used the “cuttree” R function to determine cell type specific DNA methylation programs and randomly selected 350 CpG sites from each of these clusters for visualization.

### Correlation to gene expression data

RNA-sequencing (RNA-Seq) raw counts were downloaded from the Haemopedia collection and processed using the *DESeq2* Bioconductor package [48, 49]. Genes with more than 10 reads over all cell populations were included as expressed genes. The data was subjected to variance stabilizing transformation using the *DESeq2* „vst” function. The genes associated with each CRE (based on the distance definition; see *Annotation of the MMBC array*) were extracted and used for calculation of the Pearson correlation between the cell types.

### Identification of putative associations between CREs and target genes

Differential expression from LSK cells to all differentiated downstream cell populations was calculated using the *DESeq2* Bioconductor package. Shrinkage of log2 foldchange (log2FC) values was performed using the adaptive shrinkage estimator (ashr) as implemented in the “lfcShrink” *DESeq2* function [49]. Genes with an FDR-adjusted p-value <0.01 and log2FC >2 were considered as differentially expressed genes (DEGs). Transcription start sites (TSS) of expressed genes were gathered from the Ensembl Archive Release 102 (November 2020; GRCm38.p6) and all TSSs within a distance of 1Mb to novel mdCREs were mapped. For each novel mdCRE/gene pair, the mean methylation β-value and mean vst-adjusted expression value were calculated across the replicates of each cell type. These sets of paired methylation/expression values were subjected to an association test using Pearson’s correlation and a linear model (i.e. expression of the target gene as a function of the methylation value). The correlation test p-values were false discovery rate (FDR) adjusted using the Benjamini-Hochberg method. Sites with an FDR-adjusted correlation test p-value <0.01 were considered as candidate associations. The intensity of association could be defined as the slope of the underlying linear model whereby an absolute slope >4 was required for putative novel mdCRE/gene pair.

### Analysis of HiC data

HiC data of naïve CD8+ T-cells was downloaded from GEO (GSM5017661) and analyzed using *Juicebox* [50, 51]. Associated novel mdCREs for *Thy1* were lifted to mm9 and annotated together with the *Thy1* TSS.

### Data and code availability

All statistical analyses were performed using R version 4.0.3 and the code is publicly available on GitHub (https://github.com/MaxSchoenung/MMBC). The *ggplot2* and *pheatmap* (https://github.com/raivokolde/pheatmap) R packages were used for visualization [52]. The MMBC data for hematopoietic cell types is available at the NCBI’s Gene Expression Omnibus (GEO) data repository (GSE201923). In addition, the following publicly available datasets were used for this study: Haemopedia-Mouse-RNASeq (https://www.haemosphere.org/datasets/show), LSK TWGBS (GSE146907), Hematopoiesis RRBS (https://medical-epigenomics.org/papers/broad_mirror/invivomethylation/index.html), CD8+ T-cells HiC (GSM5017661)[9, 42, 48, 50].

## DECLARATIONS

### ETHICS APPROVAL AND CONSENT TO PARTICIPATE

All mouse experiments were approved by local authorities according to German and European guidelines.

### CONSENT FOR PUBLICATION

Not applicable.

### AVAILABILITY OF DATA AND MATERIALS

The dataset generated during the current study is available in the NCBI’s Gene Expression Omnibus (GEO) data repository (GSE201923).

### AUTHORSHIP CONTRIBUTIONS

M.Schö. and D.B.L. jointly designed and coordinated the study. M.Schö., M.Hart., S.K., S.S., M.Hak., E.K., T.A., D.C., P.L. and D.B.L. performed experiments and/or analyzed and interpreted the data. M.Schö., S.K., and P.L. performed bioinformatic analysis. F.H., S.F., M.D.M., M.Schl., P.L. and D.B.L. contributed samples, and/or materials, resources and reagents. M.Schö., M.Hart. and D.B.L. wrote the first draft of the manuscript. All co-authors contributed to the final version of the manuscript.

## ACKNOWLEDGEMENTS

We thank the Microarray unit of the Genomics and Proteomics Core Facility, German Cancer Research Center (DKFZ), for providing the Illumina Mouse Methylation BeadChip arrays and related services. Additionally, we want to thank the Flow Cytometry Core Facility (DKFZ) and the Central Animal Laboratory (DKFZ) for their excellent support. We also want to thank all members of the Division of Translational Medical Oncology (DKFZ & NCT) for helpful discussions related to this study. Figure 1b was created with BioRender.com.

---

**Supplementary Figure 1.**
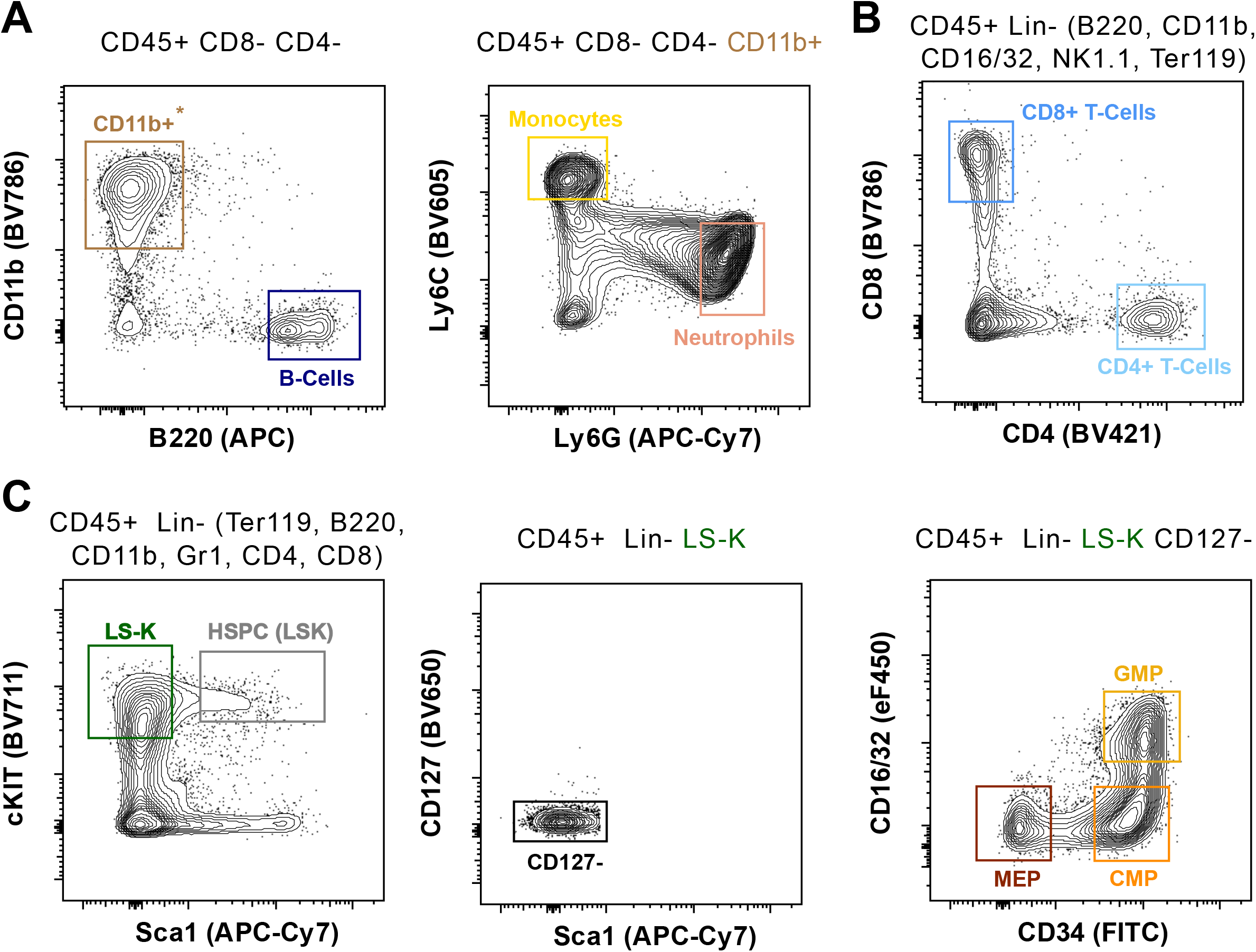
Gating strategy for the isolation of reference cell types. Flow cytometry density plots depicting the cell surface marker-based gating schemes for the isolation of hematopoietic cell types. Previous gates are listed above the respective plots. **(A)** Monocytes (CD45+ CD8- CD4- CD11b+ Ly6C^hi^ Ly6G-), Neutrophils (CD45+ CD8- CD4- CD11b+ Ly6C- Ly6G+) and B-cells (CD45+ CD8- CD4- CD11b+ B220+) were sorted without linea ge depletion. **(B)** Differentiated cells which are not from the T-cell lineage were depleted for the isolation of CD4+ T-cells (CD45+ Lin- CD4+ CD8-) and CD8+ T-cells (CD45+ Lin- CD4- CD8+). **(C)** LSKs (CD45+ Lin- Sca1+ cKIT+), MEPs (CD45+ Lin- Sca1- cKIT+ CD127- CD34- CD16/32-), CMPs (CD45+ Lin- Sca1- cKIT+ CD127- CD34+ CD16/32-) and GMPs MEPs (CD45+ Lin- Sca1- cKIT+CD127- CD34+ CD16/32+) were sorted after depletion of differentiated hematopoietic cells.

**Supplementary Figure 2.**
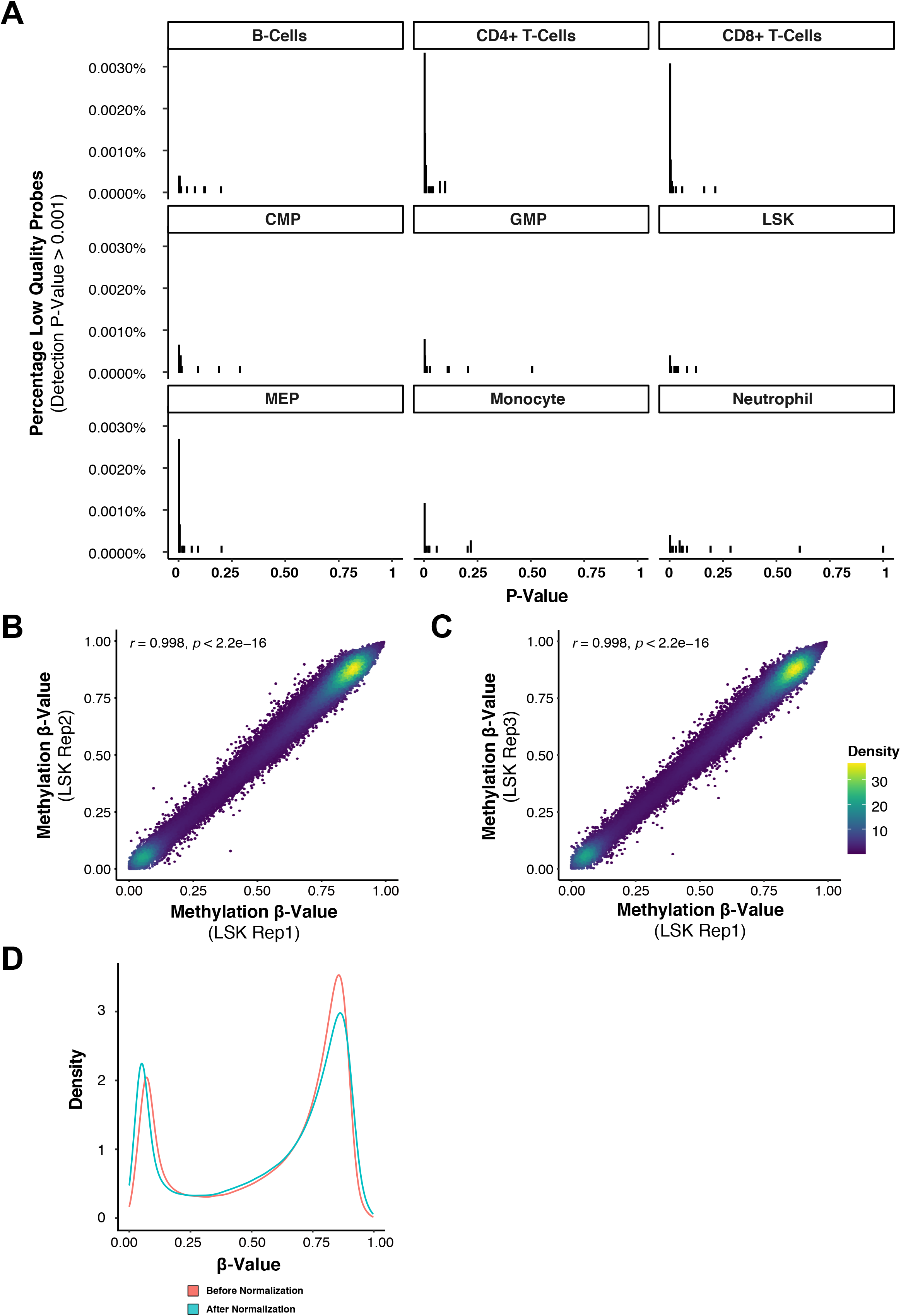
Quality control of MMBC samples. **(A)** Histograms showing the detection p-values for sites on the MMBC with a low detection rate (p-value > 0.001) per cell type. The percentage of these sites relative to all sites on the MMBC array is shown on the y-axis. **(B+C)** Scatter density plots depicting the methylation β-values of all CpG sites across the three biological replicates analyzed from LSK cells on the MMBC array. Pearson’s correlation coefficient is provided for each comparison. (**D)** Density plot showing methylation β-value distribution before and after intra-array normalization.

**Supplementary Figure 3.**
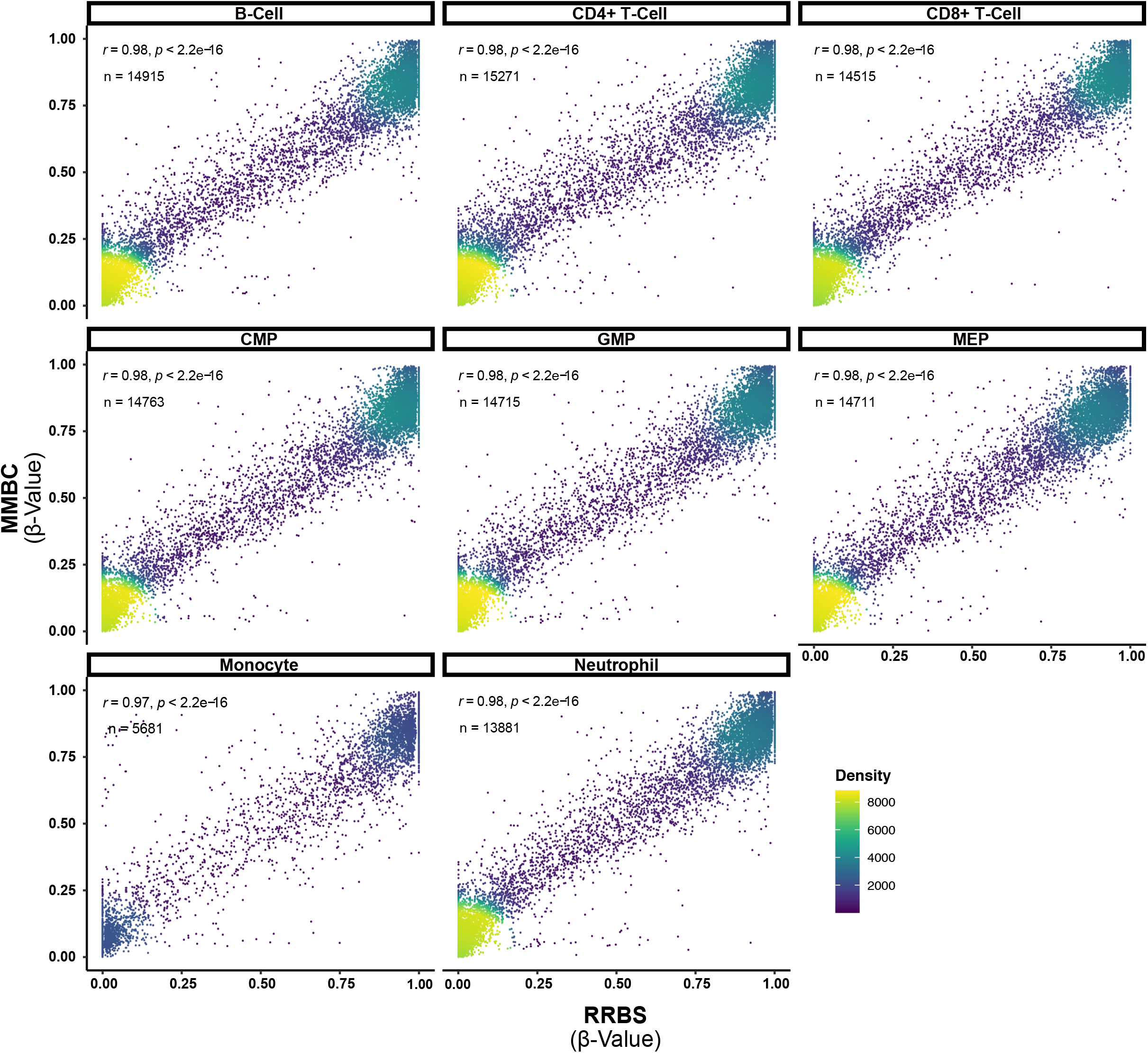
Correlation between MMBC and RRBS data. Scatter density plots showing the methylation β-values of corresponding CpG sites from RRBS (x-axis) and MMBC (y-axis) analyses per cell type. The total number of CpG sites with > 20x read coverage in RRBS is annotated together with the Pearson’s correlation coefficient.

**Supplementary Figure 4.**
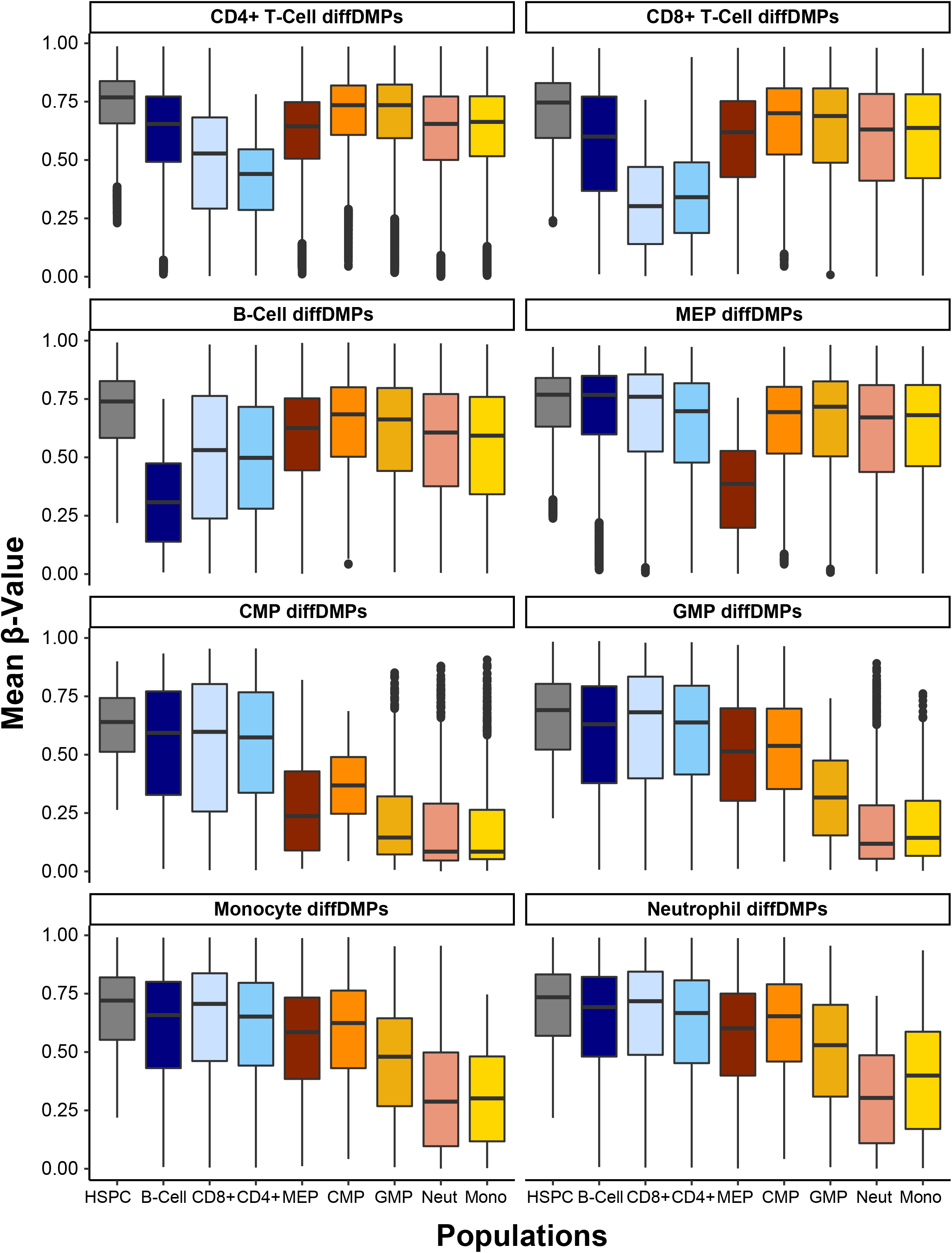
DNA methylation changes at hypomethylated diffDMPs. Boxplots showing the mean methylation β-values over the replicates for cell type specific hypomethylated diffDMPs (panel headings) per cell type.

**Supplementary Figure 5.**
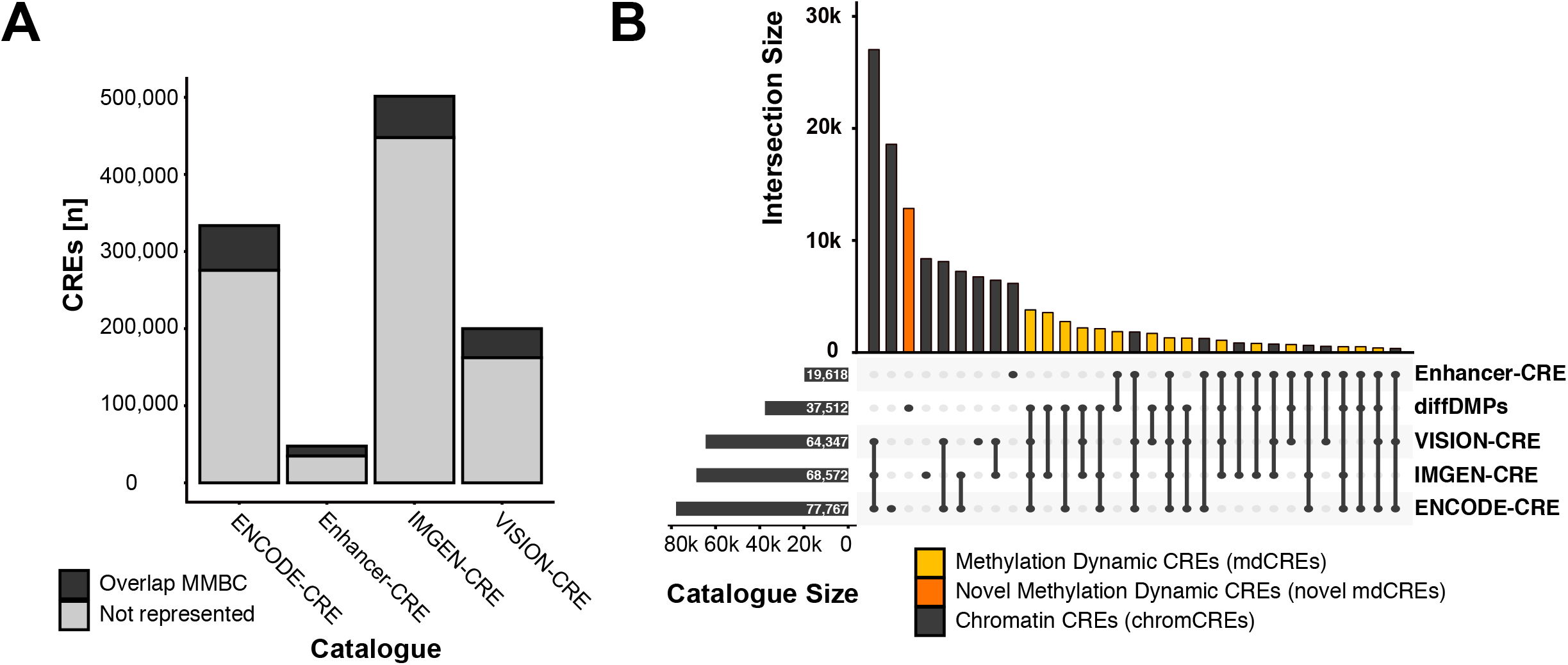
Characterization of MMBC probes overlapping with CRE catalogues. **(A)** Barplot showing the number of CREs per catalogue which overlap with the MMBC array. **(B)** Upset plot showing the mutual overlap of CRE-catalogues and diffDMPs. The intersection size is showing as column barplots and the catalogue sizes as row barplots. Chromatin CREs (chromCREs; grey), methylation dynamic CREs (mdCREs; yellow) and novel methylation dynamic CREs (novel mdCRE; orange) have been highlighted.

**Supplementary Figure 6.**
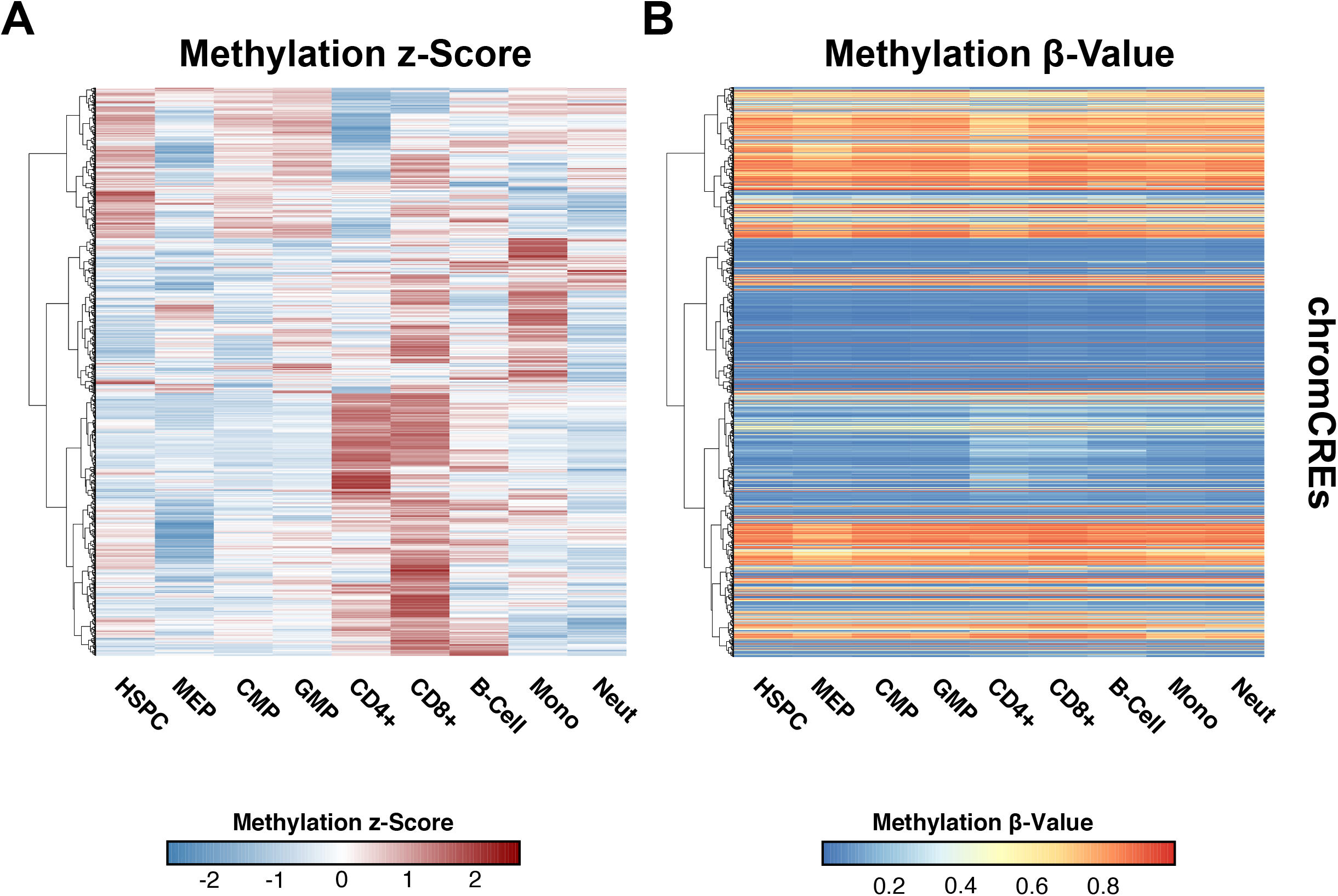
DNA methylation-based clustering of chromCREs. Methylation β-values of CpG sites from the chromCRE subset were z-score transformed and hierarchically clustered using Euclidean distance and Ward’s method. The heatmaps show methylation z-scores **(A)** and the respective β-values for 1000 randomly chosen CpG sites in the same order **(B)**.

**Supplementary Figure 7.**
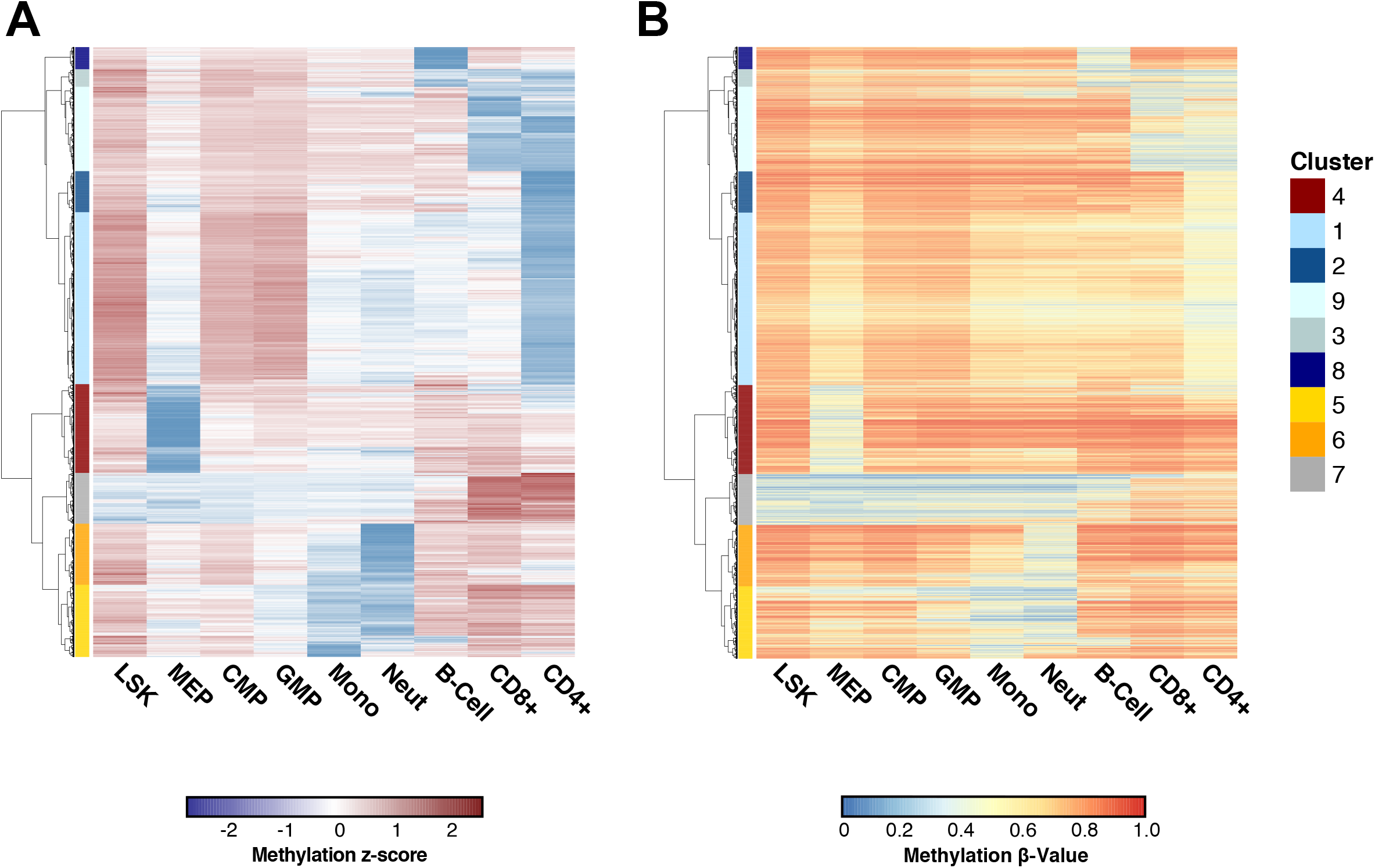
Clustering of novel mdCREs. Methylation β-values of all novel mdCREs were z-score transformed and hierarchically clustered using Euclidean distance and Ward’s method. This strategy identified nine different clusters with cell type specific DNA methylation patterns. Depicted are the z-score transformed data **(A)** and the respective absolute β-values **(B)**, which were plotted in the same order.

**Supplementary Figure 8.**
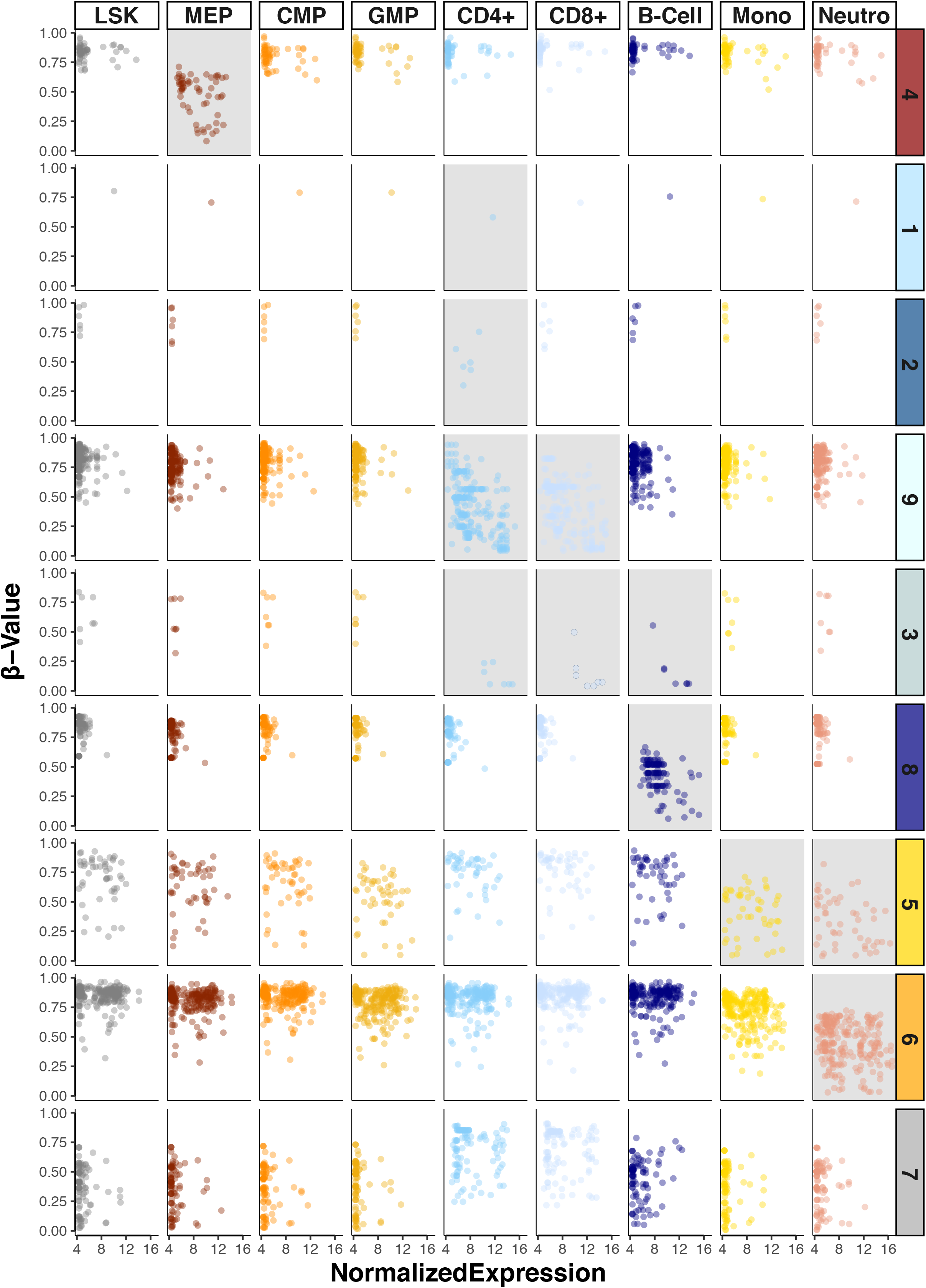
DNA methylation and gene expression dynamics for novel mdCRE-gene pairs. Methylation β-values and normalized gene expression were plotted for the identified 843 novel mdCRE-gene pairs across all cell types. The novel mdCRE were stratified by DNA methylation cluster.

